# Identifying genomic regions and candidate genes selected during the breeding of rice in Vietnam

**DOI:** 10.1101/2021.08.04.455072

**Authors:** Janet Higgins, Bruno Santos, Tran Dang Khanh, Khuat Huu Trung, Tran Duy Duong, Nguyen Thi Phuong Doai, Anthony Hall, Sarah Dyer, Le Huy Ham, Mario Caccamo, Jose De Vega

## Abstract

**Background and aims:** Vietnam harnesses a rich diversity of rice landraces adapted to a broad range of conditions, which constitute a largely untapped source of genetic diversity for the continuous improvement of rice cultivars. We previously identified a strong population structure in Vietnamese rice, which is captured in five Indica and four Japonica subpopulations, including an outlying *Indica-5* group. Here, we leveraged on that strong differentiation, and the 672 rice genomes generated, to identify genes within genomic regions putatively selected during domestication and breeding of rice in Vietnam.

**Methodology:** We identified significant distorted patterns in allele frequency (XP-CLR method) and population differentiation scores (F_ST_), resulting from differential selective pressures between native subpopulations, and compared them with QTLs previously identified by GWAS in the same panel. We particularly focused on the outlying *Indica-5* subpopulation because of its likely novelty and differential evolution.

**Results:** We identified selection signatures in each of the Vietnamese subpopulations and carried out a comprehensive annotation of the 52 regions selected in *Indica-5*, which represented 8.1% of the rice genome. We annotated the 4,576 genes in these regions, verified the overlap with QTLs identified in the same diversity panel and the comparison with a F_ST_ analysis between subpopulations, to select sixty-five candidate genes as promising breeding targets, several of which harboured alleles with non-synonymous substitutions.

**Conclusions:** Our results highlight genomic differences between traditional Vietnamese landraces, which are likely the product of adaption to multiple environmental conditions and regional culinary preferences in a very diverse country. We also verified the applicability of this genome scanning approach to identify potential regions harbouring novel loci and alleles to breed a new generation of sustainable and resilient rice.

**Key Message:** We localised regions in the rice genome selected during breeding by comparing allele frequency patterns among Vietnamese rice subpopulations. We characterised candidate genes in the Indica-5 subpopulation with breeding potential.

## Introduction

Vietnam harnesses a rich novel rice diversity due to the presence of native and traditional rice varieties adapted to its broad latitudinal range, diversity of ecosystems and regional food preferences (Fukuoka et al. 2003). This diversity constitutes a largely untapped and highly valuable genetic resource for local and international breeding programs (Khanh et al. 2021). Vietnamese rice shows a strong population structure, which is captured within five Indica and four Japonica subpopulations that we have recently described (Higgins et al. 2021). These subpopulations were characterised in relation to the fifteen subpopulations of Asian rice described by the rice 3,000 rice genomes project (3K RGP) (Zhou et al. 2020). Among these nine populations described in Vietnam, the *Indica-5* (I5) subpopulation is an outlier and is expanded in Vietnam and, therefore, a potential source of novel variation compared to the wider Asian diversity.

Genetic variation and differentiation are influenced by natural processes, such as adaption and random drift, as well as conscious breeding selection and unconscious selection, due to the agricultural practices of local farmers. Selection causes detectable changes in allele frequencies at the selected sites and their flanking regions. By modelling differences in allele frequency in close loci between neutrality and selection scenarios, the cross-population composite likelihood ratio test (XP-CLR) can detect selective sweeps (Chen, Patterson, and Reich 2010). Any distorted pattern in allele frequency in contiguous SNP sites would have occurred too quickly (speed of change is assessed over expanding windows based on the length of the affected region) to be explained by random drift, so allowing the identification of selected regions. XP-CLR has been used to identify regions of selection associated with domestication and improvement in a wide range of crops, such as apple (Duan et al. 2017), soybean (Zhou et al. 2015), cucumber (Qi et al. 2013) and wheat (Joukhadar et al. 2019); and has been increasingly used in rice. Lyu et al.(2014) identified a list of differentiated genes that may account for the phenotypic and physiological differences between upland and irrigated rice. Xie et al (2015) compared Indica semi-dwarf modern bred varieties (IndII) with taller Chinese landraces (IndI) to identify signatures of rice improvement and detected 200 regions spanning 7.8% of the genome. Meyer et al. (2016) identified genomic regions associated with adaptive differentiation between *O. glaberrima* populations in Africa. He et al. (2017) tested for positive selection between weedy and landrace rice using five different approaches. Cui et al. (2019) identified potential selective sweeps in both Indica and Japonica genomes showing that there were multiple loci responding to selection and that loci associated with agronomic traits were particularly targeted by selection. Lin et al. (2020) used XP-CLR to demonstrate how introgressed regions were selected through hybrid rice breeding. While these studies were trying to answer difference questions, all used XP-CLR to detect selected regions. In addition, many of the studies used other metrics, such as the fixation index (F_ST_), to verify selected regions.

Here, we identified regions in the rice genome which have been selected by human breeding by leveraging the strong population structure among Vietnamese-native rice varieties and landraces, which has resulted from its diverse geography and agronomic practices. Unravelling the genomic differences and identifying regions selected between these nine subpopulations is the first step towards understanding their breeding potential. To assess the potential of our approach, we later focused on the outlying *indica-5* (I5) subpopulation to identify candidate loci for breeding targets, since this subpopulation constitutes a gene-pool not used in rice improvement. To assess the putative role of these selected regions and whether these selected regions may contain loci that potentially could control agronomic traits, we looked for overlaps with previously mapped QTLs in the same diversity panel, and regions enriched in gene ontology (GO) terms. QTLs have been described for a range of agronomic traits using the complete set of 672 native rice accessions (Higgins et al. 2021), while a subset of 182 of these traditional Vietnamese accessions (Phung 2014) were used for genome-wide phenotype-genotype association studies (GWAS) relating to root development (Phung et al. 2016), panicle architecture (Ta et al. 2018), drought tolerance (Hoang, Van Dinh, et al. 2019), leaf development (Hoang, Gantet, et al. 2019), Jasmonate regulation (To et al. 2019) and phosphate starvation and efficiency (Mai et al. 2020; To et al. 2020). Finally, we studied alleles with non-synonymous substitutions in candidate genes in selected regions of the outlying and highly selected I5 subpopulation.

## Materials and Methods

### Sequencing and SNP calling and annotation

A set of 616 Vietnamese samples were sequenced by us as described previously (Higgins et al. 2021). We added 56 samples from the “3000 Rice Genomes Project” (3K RGP), which originated from Vietnam, to give a total of 672 samples. These samples were genotyped as described previously (Higgins et al. 2021) to obtain a set of imputed SNPs which were filtered to remove SNPs with excess heterozygosity, followed by filtering for a minimal allele frequency of 5%. We utilized a set of 2,027,294 SNPs for the 426 Indica samples and 1,125,716 SNPs for the 211 Japonica samples. Detailed information for each sample is available in Higgins et al. (2021). A summary of the number and source of each subpopulation is available in Table 1 (47 Indica samples and 9 Japonica samples native to Vietnam were obtained from the 3K RGP project). The putative effects of the bi-allelic SNPs (low, medium and high effects) on the genome were determined using SnpEff (Cingolani et al. 2012) as detailed in Higgins et al. (2021) and the pre-built release 7.0 annotation from the Rice Genome Annotation Project (http://rice.plantbiology.msu.edu/).

**Table 1.**
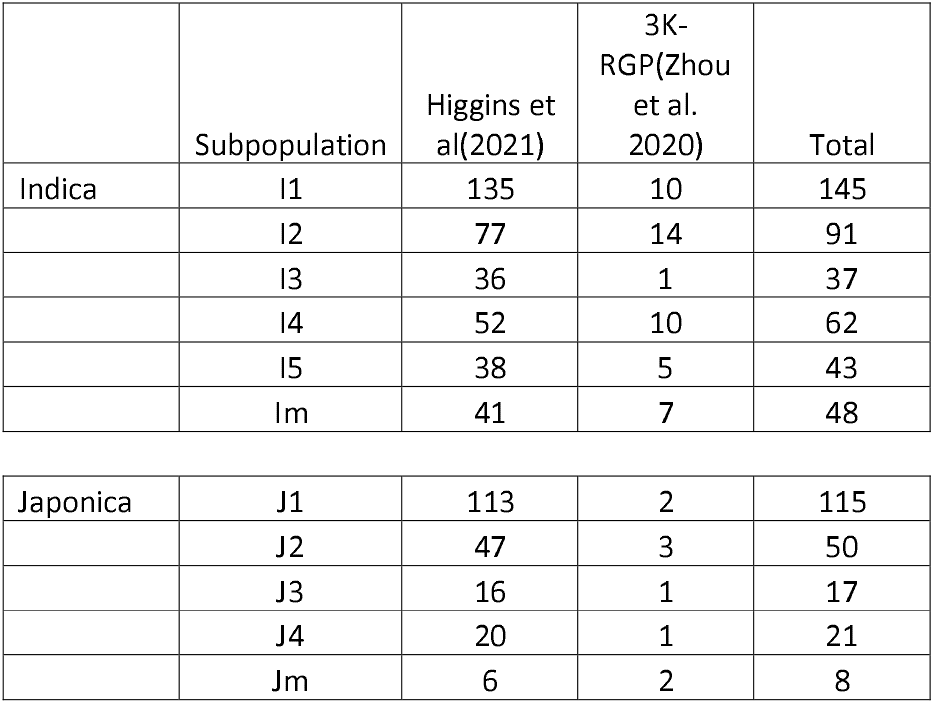
Summary of the 426 Indica and 211 Japonica samples. The complete dataset of 672 samples includes 35 admixed samples, full sample detail in Higgins et al.(2021)

### Identification of selective sweeps using XP-CLR

Selective sweeps across the genome were identified using XP-CLR (Chen, Patterson, and Reich 2010), a method based on modelling the likelihood of multilocus allele frequency differentiation between two populations. An updated version of the original code was used (https://github.com/hardingnj/xpclr). We used 100 kbps sliding windows with a step size of 10 kbps and the default option of a maximum of 200 SNPs in any window. XP-CLR was run comparing the five Indica subpopulations to each other and the four Japonica subpopulations to each other. Selected regions were extracted using the XP-CLR score for each 100 kbps window as follows: 200 kbps centromeric regions were removed. The mean and 99th percentile of the XP-CLR scores were calculated for each comparison between one subpopulation against the remaining ones (e.g. I5 vs I1, I2, I3, I4). The mean 99^th^ percentile was used to define the cut-off level for selection in that subpopulation. 100 kbps regions with an XP-CLR score higher than the cut-off were extracted and contiguous regions were merged using BEDTools v2.26.0 (Quinlan and Hall 2010) specifying a maximum distance between regions of 100 kbps. Regions shorter than 80 kbps were removed to give a final set of putatively selected regions for each comparison. Putative regions observed selected in at least two comparisons for Japonica subpopulations, or three comparisons for Indica subpopulations, were merged to obtain a final set of selected regions for each subpopulation. BEDTools map was used for finding any overlap of selected regions with QTLs. QTL regions using the same, or a subset of, the samples were previously identified by reviewing the literature. Genes lying within the selected regions were extracted and checked for enrichment in Protein Domain and Pathway using a maximum Bonferroni FDR value of 0.05 in PhytoMine (https://phytozome.jgi.doe.gov/), a service implemented within Phytozome (Goodstein et al. 2012).

### Calculating F_ST_

We calculated F_ST_ per SNP between the 43 samples in the I5 subpopulation and the 190 samples in the I2, I3 and I4 subpopulations with VCFtools using the “weir-fst-pop” option, which calculates F_ST_ according to the method of Weir and Cockerham (Weir and Cockerham 1984). Sites which are homozygous between these populations were removed, and negative values were changed to zero. The mean F_ST_ was calculated per gene and per specified region.

### Enrichment analysis of GO terms in selected regions

The enrichment analysis was done with the library topGO (Alexa A 2010) in R, using as inputs the lists of genes in each selected region, and the functional annotation of the rice genome (Rice MSU7.0) from agriGO (http://bioinfo.cau.edu.cn/agriGO). The method in topGO compared the genes observed in each selected region annotated with a given GO term with the expected number of genes annotated with that term in the whole transcriptome. The statistical test was a F-Fisher test (FDR□<□0.05) with the “weight01” algorithm in topGO. The “weight01” algorithm resolves the relations between related GO ontology terms at different levels. The number of genes and selected regions that were enriched terms for the different subpopulations were plotted using ggplot2 (Wickham 2016).

### Results Identification of selective sweeps among Vietnamese subpopulations

To identify genomic regions that have been selected during the breeding of rice in Vietnam, we searched for genomic regions with distorted patterns of allele frequency that cannot be explained by random drift using XP-CLR (Chen, Patterson, and Reich 2010). We used our previously described dataset of 672 genomes from Vietnamese-native landraces and varieties (Table 1), which have been divided into nine subpopulations (Higgins 2021). We compared all the five Indica subpopulations to each other and all the four Japonica subpopulations to each other. Firstly, we obtained the mean XP-CLR score over the whole genome, as summarised in Table 2, with the reciprocal differences in the comparisons between a pair of subpopulations in Supplementary Table S1. Among the Japonica subpopulations, the J4 subpopulation had consistently the highest selection scores especially against the J1 subpopulation. Among the Indica subpopulations, the I1 subpopulation had consistently the lowest selection scores. The I5 subpopulation had the highest selection scores except in comparison with the I3 subpopulation. We calculated the 99^th^ percentile for each comparison between a pair of subpopulations and used the mean value for each subpopulation as a cut-off to identify selected regions (detailed in Supplementary Table S2 and summarised in Table 3). We merged selected regions within 100kb of each other, so the final set of selected regions for each comparison were of variable length. Selected regions covered a higher proportion of the genome where the XP-CLR score was higher. The regions selected in the comparisons between a pair of subpopulations were plotted along each chromosome for the Indica subpopulations (Supplementary Fig. S1) and the Japonica subpopulations (Supplementary Fig. S2). In order to define a final set of selected regions in a given subpopulation, we retained and merged regions selected in at least three comparisons between that subpopulation and any other subpopulation in the case of the Indica ones, or in at least two comparisons in the case of the Japonica subpopulations. This procedure is described in detail for the I5 subpopulation in a subsequent section. The final set of selected regions in each subpopulation were plotted along each of the rice chromosomes in Fig. 1A and 1B for the Indica and Japonica subtypes, respectively. The selected regions ranged from 98,583 to 2,787,579 bases for the Japonica subpopulations, and from 106,844 to 2,309,615 bases for the Indica subpopulations. We observed slightly different patterns in length variation per subtype and subpopulation (Supplementary Fig. S3). Overall, the Japonica subpopulations had fewer selected regions, which represented from 3.7 to 4.9% of the genome, while Indica subpopulations ranged from 5.3 to 8.1% of the genome. Gene lists for the selected regions are available in Supplementary Table S3. The Japonica subtypes had a higher proportion of long selected regions. These regions were confined to specific areas of the genome and absent from large chromosome regions. All four Japonica subpopulations were selected on the long arm of chromosome 2 and in both flanks of the centromeric region of chromosome 4. The selected regions in the Indica subpopulations were spread throughout the genome and very variable in length. We particularly observed a high proportion of shorter than average selected regions and a lower proportion of longer than average selected regions in the I1 subpopulation. The I5 subpopulation stands out as having the highest proportion of the genome under selection, overlapping with the other landrace subpopulations (I2, I3 and I4) on the short arm of chromosome 1 and the long arm of chromosome 9. However, selected regions in I5 were absent on the long arm of chromosome 4, where all other landrace subpopulations overlapped with the elite I1 subpopulation.

**Table 2.** Whole-genome XP-CLR selection scores. Mean XP-CLR score across the whole genome for each comparison between the four Japonica subpopulations and the five Indica subpopulations. Reciprocal comparisons shown in Supplementary Table S1.

|  | SCORE | J1 | J2 | J3 | J4 |
| --- | --- | --- | --- | --- | --- |
| Selected | J1 |  | 17.8 | 7.6 | 6.1 |
|  | J2 | 19.5 |  | 21.6 | 6.6 |
|  | J3 | 24.4 | 17.9 |  | 5.9 |
|  | J4 | 46.1 | 17.5 | 17.9 |  |

|  | SCORE | I1 | I2 | I3 | I4 | I5 |
| --- | --- | --- | --- | --- | --- | --- |
| Selected | I1 |  | 8.5 | 4.0 | 8.2 | 9.8 |
|  | I2 | 28.8 |  | 7.0 | 15.7 | 17.3 |
|  | I3 | 40.2 | 24.5 |  | 23.2 | 23.7 |
|  | I4 | 34.1 | 21.5 | 7.4 |  | 18.6 |
|  | I5 | 63.6 | 44.6 | 18.0 | 39.2 |  |

**Table 3.** XP-CLR scores and summary on the regions under selection in each subpopulation. Individual comparisons are shown in Supplementary Table S2

|  | mean<br>xp_clr<br>score | cutoff (99<br>percentile) | filtered<br>regions<br>>80000bp | mean<br>length | total<br>length | % of<br>genome<br>373,245,519<br>bp | number of<br>genes |
| --- | --- | --- | --- | --- | --- | --- | --- |
| J1 | 10.5 | 136 | 28 | 576,707 | 16,147,785 | 4.3 | 2427 |
| J2 | 25.9 | 256 | 23 | 726,689 | 16,713,841 | 4.5 | 2439 |
| J3 | 16.1 | 228 | 24 | 577,089 | 13,850,139 | 3.7 | 2007 |
| J4 | 27.1 | 297 | 25 | 731,341 | 18,283,522 | 4.9 | 2643 |
| I1 | 7.6 | 161 | 44 | 453,570 | 19,957,065 | 5.3 | 3077 |
| I2 | 17.2 | 275 | 41 | 550,836 | 22,584,270 | 6.1 | 3346 |
| I3 | 27.9 | 401 | 42 | 474,009 | 19,908,387 | 5.3 | 2993 |
| I4 | 20.4 | 306 | 38 | 619,404 | 23,537,343 | 6.3 | 3465 |
| I5 | 41.4 | 440 | 52 | 583,706 | 30,352,734 | 8.1 | 4576 |

**Fig. 1.**
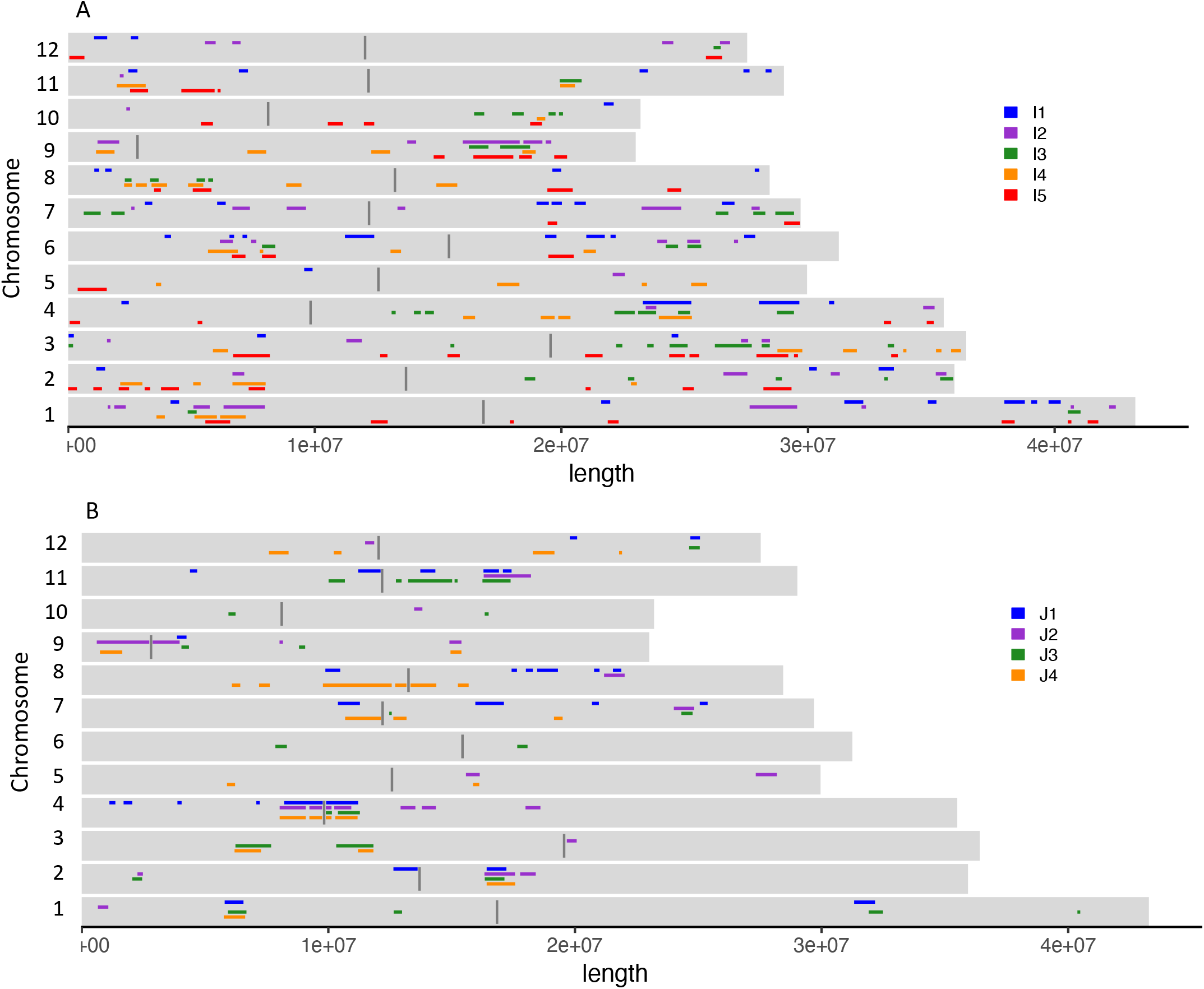
XP-CLR scores and regions under selection. (a) Selected regions for the five Indica subpopulations covering 5.4%, 6.1%, 5.3%, 6.3% and 8.1% of the genome for I1, I2, I3, I4 and I5 respectively. Centromeric regions are shown as 100 kb regions in dark grey. (b) Selected region for the four Japonica subpopulations covering 4.3%, 4.5%, 3.7% and 4.9% of the genome for J1, J2, J3 and J4 respectively.

### Putative roles of the regions under selection

We looked for the overlap of the selected regions with sets of QTLs previously reported in the literature (Supplementary Tables S4 and S5); 21 QTLs for basic plant and seed architecture traits identified using the same complete set of Vietnamese rice samples (Higgins el al.2021); and 88 QTLs associated with root development traits (Phung et al. 2016), 29 QTLs for panicle morphological traits (Ta et al. 2018), 17 QTLs for tolerance to water deficit (Hoang, Van Dinh, et al. 2019), 13 QTLs for leaf mass traits (Hoang, Gantet, et al. 2019), 25 QTLs for growth mediated by jasmonate (To et al. 2019), 21 QTLs for phosphate starvation (Mai et al. 2020) and 18 QTLs for phosphate efficiency (To et al. 2020) reported for a subset of 180 samples of the whole dataset. The selected regions in the Japonica subpopulations had overlaps with all the QTLs sets, except QTLs associated with growth regulation by jasmonate (Supplementary Table S5). The region on chromosome 2 that was selected in all Japonica subpopulations overlapped with a QTL for grain length (2_GL) and two related QTLs for panicle morphology, secondary branch number (SBN) and spikelet number (SpN). These QTLs collocate with osa-MIR437 (Ta et al. 2018), a monocot preferential miRNA that targets LOC_Os02g18080 (https://rapdb.dna.affrc.go.jp). J2 and J4 lowland varieties were both selected on the long arm of chromosome 5 and at the start of chromosome 9. The region on chromosome 5 overlaps with a QTL for drought sensitivity observed after 4 weeks of drought stress (q4_Score4). The selected region on chromosome 9 overlaps with a QTL for rachis length (RL), which is associated with the size of the panicle, a key component of yield. The region towards the end of chromosome 11, which was selected in J1, J2 and J3, overlaps with qRTW11.19 as well as several QTLs associated with root traits: Rq13_J_TIL, Rq29_J_DEPTH, Rq30_J_DEPTH, Rq46_F_NCR, Rq63_J_THK. The selected regions in the Indica subtypes overlapped with all the QTL sets (Supplementary Table S4). Most overlaps that occurred in more than one subpopulation were also observed in the I5 subpopulation, so are discussed in the next section. In addition, the region on the long arm of chromosome 11, which is selected in both I3 and I4, overlaps with QTLs for drought sensitivity (Tq17 Score4), rachis length (QTL25 RL) and response to jasmonate (qSHL5).

The total number of genes within the selected regions are shown in Table 2. For the Japonica subtypes, the number of genes ranged from 2,007 genes within the selected regions of the J3 subpopulation to 2,643 genes within the selected regions of the J4 subpopulation. For the Indica subtypes, the number of genes ranged from 2,993 to 3,465 in the I1 to I4 subpopulations, whilst the I5 subpopulation had 4,576 genes within 52 selected regions (gene listed in Supplementary Table S3). The overlap between genes selected in each subpopulation showed that around half of the genes selected in a subpopulation were unique to that subpopulation (Supplementary Fig. S4). No genes were selected in all subpopulations, but 230 genes were selected in all four Japonica subpopulations, and 44 genes were selected in all the Indica landrace subpopulations I2 to I5.

An enrichment analysis of the GO terms enriched in each selected region was obtained by comparing the annotations in each selected region with the whole genome annotation, as background (Supplementary Table S6). The number of genes associated with enriched terms in different regions from the same subpopulation were added up and plotted (Fig. 2). A large proportion of genes in selected regions were associated with the same biological functions in the different Indica subpopulations, e.g. Lipid and protein metabolic process, or “Biosynthetic process”. However, we also evidenced specific selections in particular subpopulations, such as “Photosynthesis” genes in I5 and J1; biotic response genes in I2, I5 and J1; abiotic response genes in I1 and I5; and “flower development” genes in I2. Selected regions were more clearly associated to specific GO terms in the Indica subpopulations than in the Japonica ones. The enrichment of GO terms was not correlated with the total number of genes or genome length in each subpopulation (Supplementary Table S2).

**Fig. 2.**
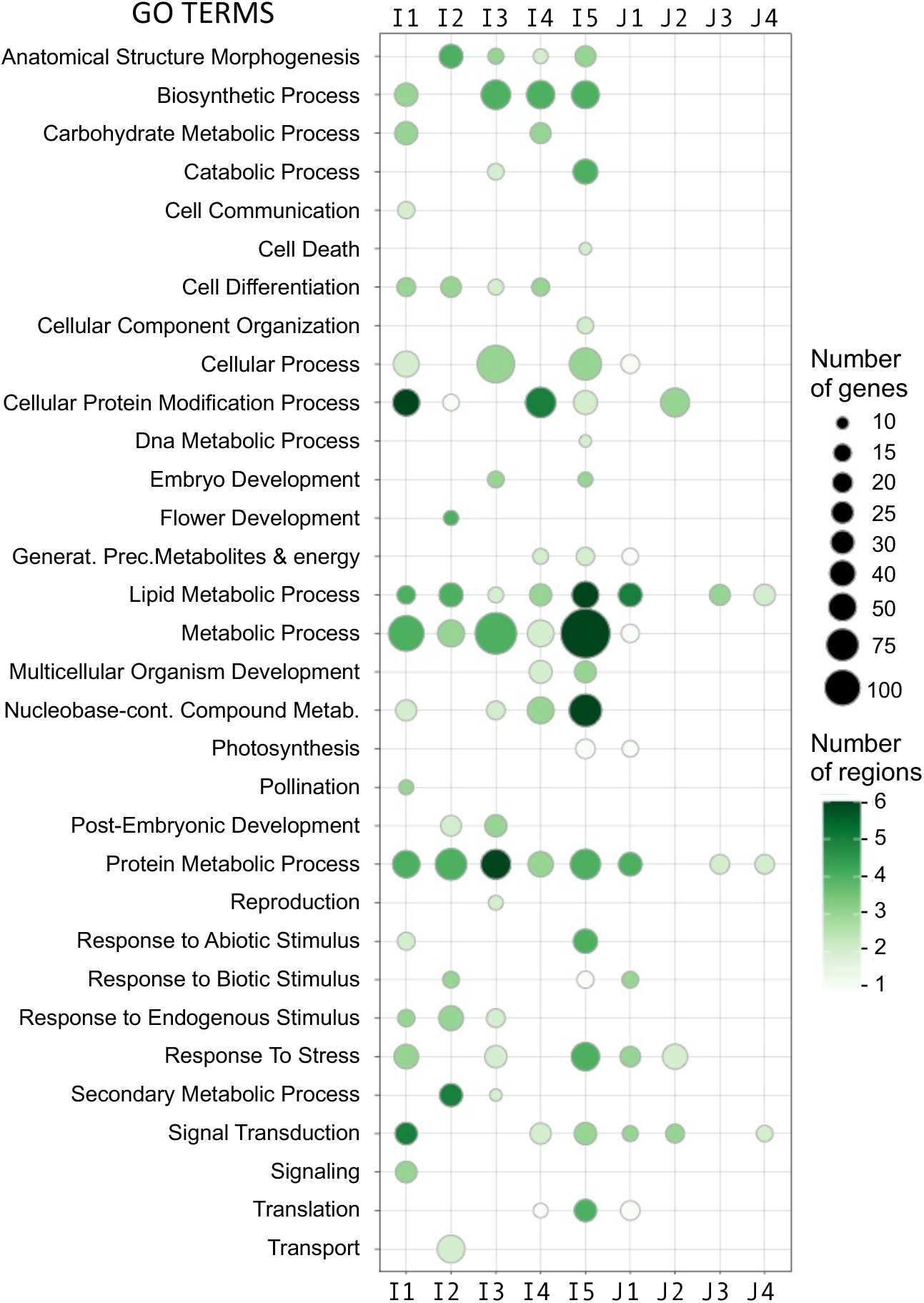
Gene Ontology overrepresentation.

### Selected regions in the outlying Indica-5 (I5) subpopulation

Overall, the I5 subpopulation had the highest XP-CLR selection scores, this is reflected in I5 having the greatest number of selected regions covering the highest proportion of the genome. I5 is an outlier subpopulation, which contains a gene-pool not present the modern bred improved varieties that comprise subpopulation I1 (Higgins et al 2021). The XP-CLR score of the I5 subpopulation compared to the other four Indica subpopulations in 100 kbps windows is shown in Fig. 3. A cut-off XP-CLR score of 440 was used to defined selected regions in I5; I5 vs I1 produced 207 regions with a mean length of 267 kbp (14.8% of the genome); I5 vs I2 produced 120 regions with a mean length of 204 kbp (6.57% of the genome); I5 vs I3 produced 14 regions with a mean length of 162 kbp (0.61% of the genome); I5 vs I4 produced 122 regions with a mean length of 122 kbp (6.02% of the genome). Regions selected in three or more subpopulations were merged to give 52 selected regions in I5, the regions are listed in Supplementary Table S7 and the functional annotation of each region detailed in Supplementary Table S8. These regions had a mean length of 584 kbp, covered 30 Mbp, which represents 8.13% of the rice genome, and contained 4,576 genes (Supplementary Table S9). To gain further information on the uniqueness of these 52 regions selected in I5, we calculated the F_ST_ per SNP between the 43 samples in the I5 subpopulation and the 190 samples in the landrace subpopulations, I2, I3 and I4. The variation of F_ST_ and diversity along each chromosome are shown in Fig. 3e and 3f. Both F_ST_ and diversity varied widely along the genome and did not show the clear peaks seen in the XP-CLR score, but peaks can be seen in F_ST_ pattern coinciding with XP-CLR peaks. This is clearest on chromosome 12 where F_ST_ and XP-CLR score showed a similar pattern and the diversity scores showing the opposite pattern. Our aim was to localise regions in the genome with both high F_ST_ between the I5 subpopulation compared to the other Vietnamese subpopulations and low diversity in the I5 subpopulation. High F_ST_ but low diversity would be expected in recently selected regions, as can be seen on chromosome 10. Chromosome 3 also showed this pattern and contained a large number of selected regions. The mean F_ST_ per gene for the 4,576 genes selected in I5 is listed in Supplementary Table S10, and the mean F_ST_ per selected region is shown in Supplementary Table S7. The 1,983,066 heterozygous SNPs in subpopulations I2, I3, I4 and I5 had a mean F_ST_ of 0.185, and this mean value increased to 0.305 for the subset of 177,874 SNPs within the I5 selected regions.

**Fig. 3.**
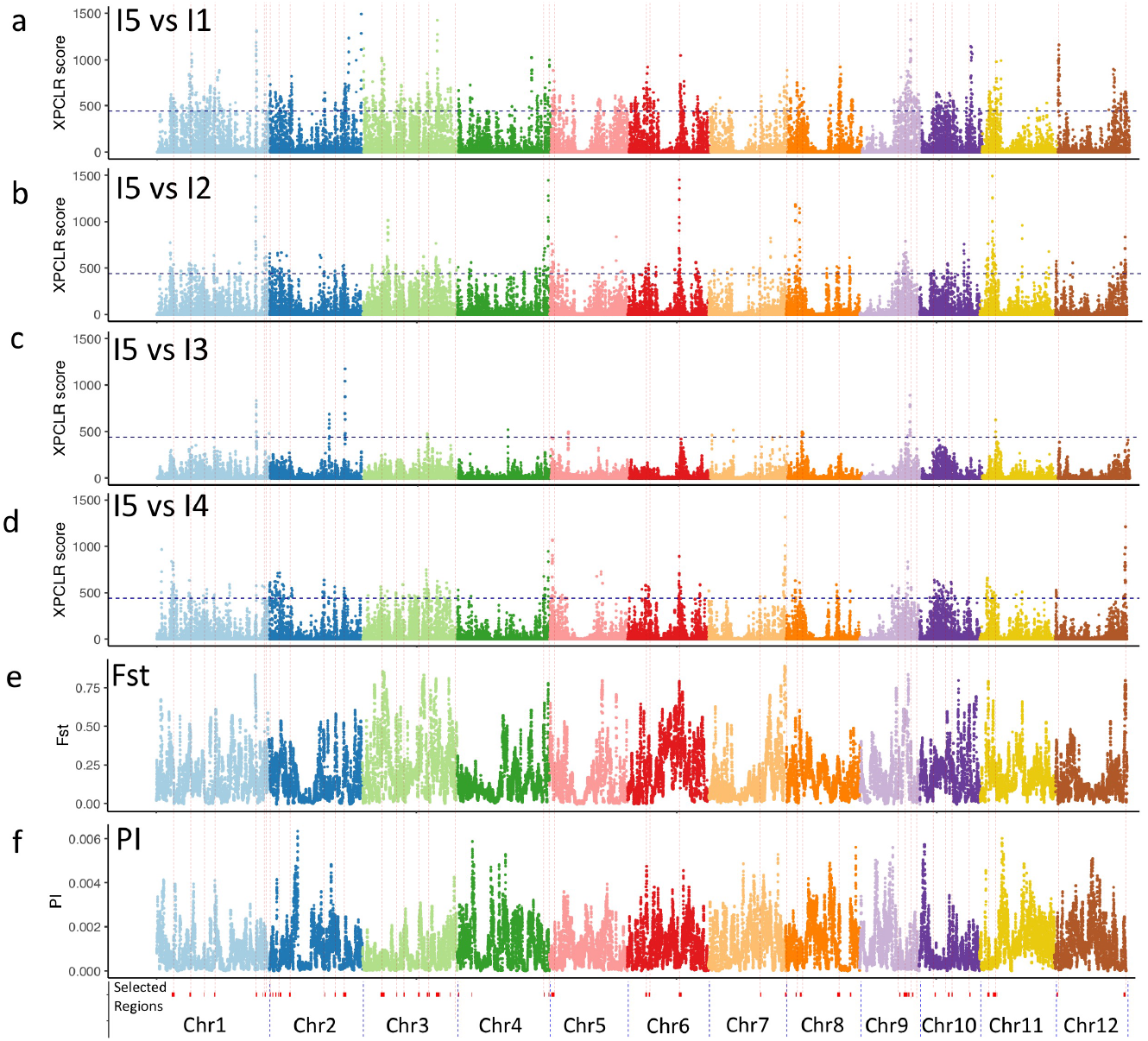
Selection sweeps in the Indica I5 subpopulation compared to the other Vietnamese subpopulations. XP-CLR scores in 100,000 bp sliding windows are plotted along the 12 chromosomes, showing selection in the I5 subpopulation compared to (a) I2, (b) I2, (c) I3, (d) I4. The horizontal dashed line indicates the threshold XP-CLR score of 440 for determining selected regions. (e) F_ST_ in 100,000 bp sliding windows for the 43 samples in the I5 subpopulation compared to the 190 samples in the I2, I3 and I4 subpopulations. (f) Whole genome genetic diversity (Π) in 100,000 bp sliding windows for the 43 samples in the I5 subpopulation. The vertical lines show the position of the 52 selected regions.

The overlap of the 52 selected regions in the I5 subpopulation with the eight sets of QTLs is shown in Fig. 4. Fourteen regions showed significant overlaps, these were shaded in Fig. 4 and listed in Table 4, detailing the individual QTLs in Supplementary Table S11. Region ‘a’ was selected in I2, I4, I5, J1, J3 and J4, and coincided with a QTL for spikelet number (SpN) and two QTLs for root traits (q73, q84) where the following candidate genes LOC_Os01g10900, LOC_Os01g11010 and LOC_Os01g11860 were located. Region ‘b’ was selected in I1 and I5, and coincided with a QTL for Leaf weight (FW_TW) and a QTL for relative phosphate uptake (qRPUpE1.5) where the candidate gene LOC_Os01g66070 was located. Region ‘c’ was selected in I3 and I5 and overlapped a QTL for response of root length to jasmonate (qRTL1), an overlap of 37 candidate genes, including the transcription factor *OsBLR1* (LOC_Os02g47660), which regulates leaf angle in rice via brassinosteroid signalling (Wang et al. 2020). Region ‘d’ partially overlapped a QTL for panicle length (9_PL) and a region selected during recent domestication by farmers in China (Cui et al. 2019). Region ‘e’ fully overlapped a QTL for grain length (12_GL). Region ‘f’ was selected in I3, I4 and I5 and coincided with a QTL for leaf length (Rq2), which is only found in the Japonica subtype. Both regions ‘e’ and ‘f’ overlapped with two large regions selected during recent domestication by farmers in China. Gene *SSIIa* (LOC_Os06g12450) and SDL/RNRS1 (LOC_Os06g14620) fall within this region. SSIIa is required for the edible quality of rice and plays an important role in grain starch synthesis (Zhang et al. 2011). SDL/RNRS1 (LOC_Os06g14620) encodes the small subunit of ribonucleotide reductase, which is required for chlorophyll synthesis and plant growth development (Qin et al. 2017). Region ‘g’ was selected in I1 and I5, and coincided with a QTL for panicle length (14_PL) and a QTL for maximum root length (Rq35). Region ‘h’ was selected in J4, I1 and I5, and coincided with a QTL for relative water content (Tq7) observed after 3 weeks of drought stress. Region ‘i’ was selected in I3 and I5, and coincided with a QTL for root depth (Rq25). Region ‘j’ was selected in I3, I4 and I5, and overlapped with a QTL for response of shoot length to jasmonate (qSHL4) and 4 candidate genes, including a G2-like transcription factor (LOC_Os08g06370). Region ‘k’ was selected in I3, I4 and I5, and coincided with two QTLs for panicle traits, primary branch number (PBN) and primary branch average length (PBL) that include the gene Auxin Response factor,*OsPILS2* (LOC_Os08g09190). Region ‘l’ coincided with a QTL for the number of crown roots in response to Pi deficiency (qNCR8.13), which contained two candidate genes, *OsPP2C66* (LOC_Os08g39100) encoding PHOSPHATASE 2C, and transcription factor *OsWKKY30* (LOC_Os08g38990). Region ‘m’ was selected in I1, I4 and I5, and coincided with four QTLs related to response of plants to drought (Tq12), which contained *OsbZIP80* (LOC_Os11g05640), a transcription factor involved in dehydration stress response (Nijhawan et al. 2008). Region ‘n’ was selected in J1 and I5, and coincided with a QTL for number of crown roots (Rq43).

**Fig. 4.**
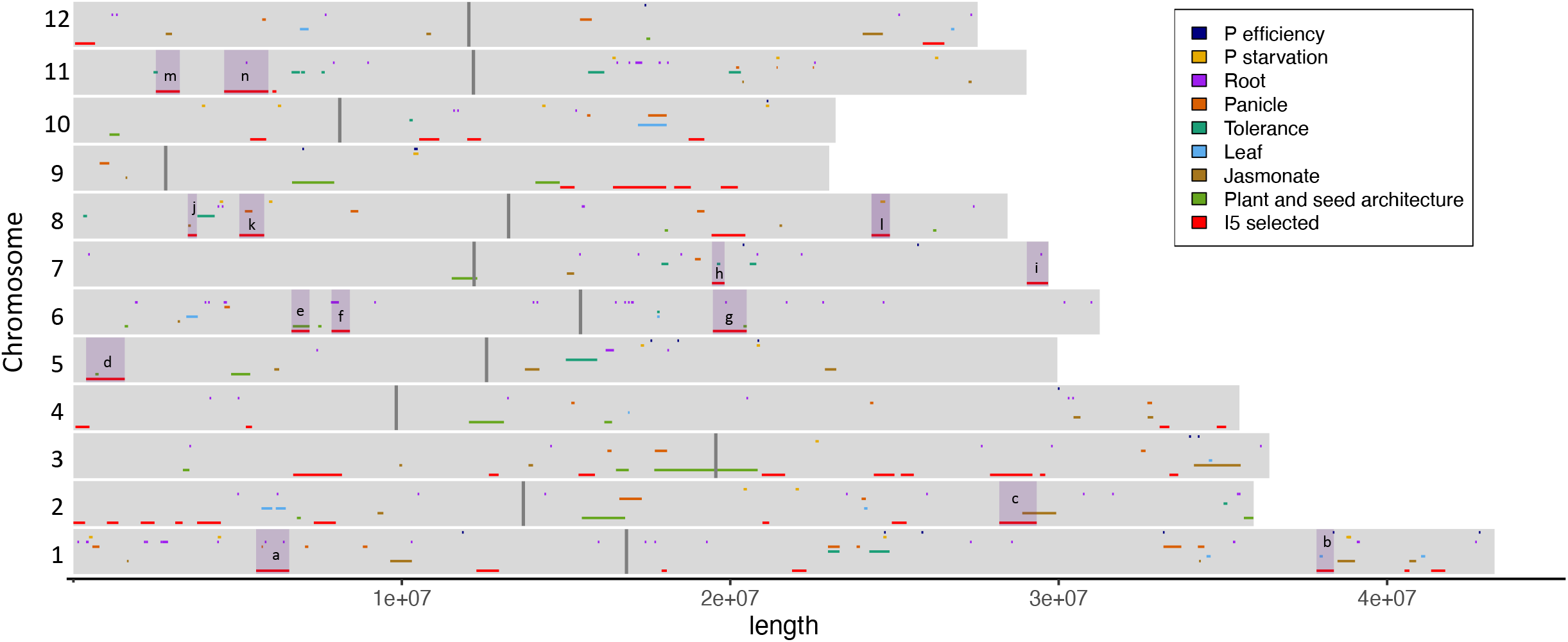
Vietnamese QTLs and their overlap with selected regions in the I5 subpopulation. QTLs from 8 published studies (Phung et al. 2016; Ta et al. 2018; To et al. 2019; Hoang, Gantet, et al. 2019; Hoang, Van Dinh, et al. 2019; Mai et al. 2020; To et al. 2020) (Higgins et al.) are plotted along each chromosome together with the 52 regions selected in the I5 subpopulation. The fourteen selected regions which overlap with at least one QTL are highlighted, the letters refer to the details shown in Table 2

**Table 4:**
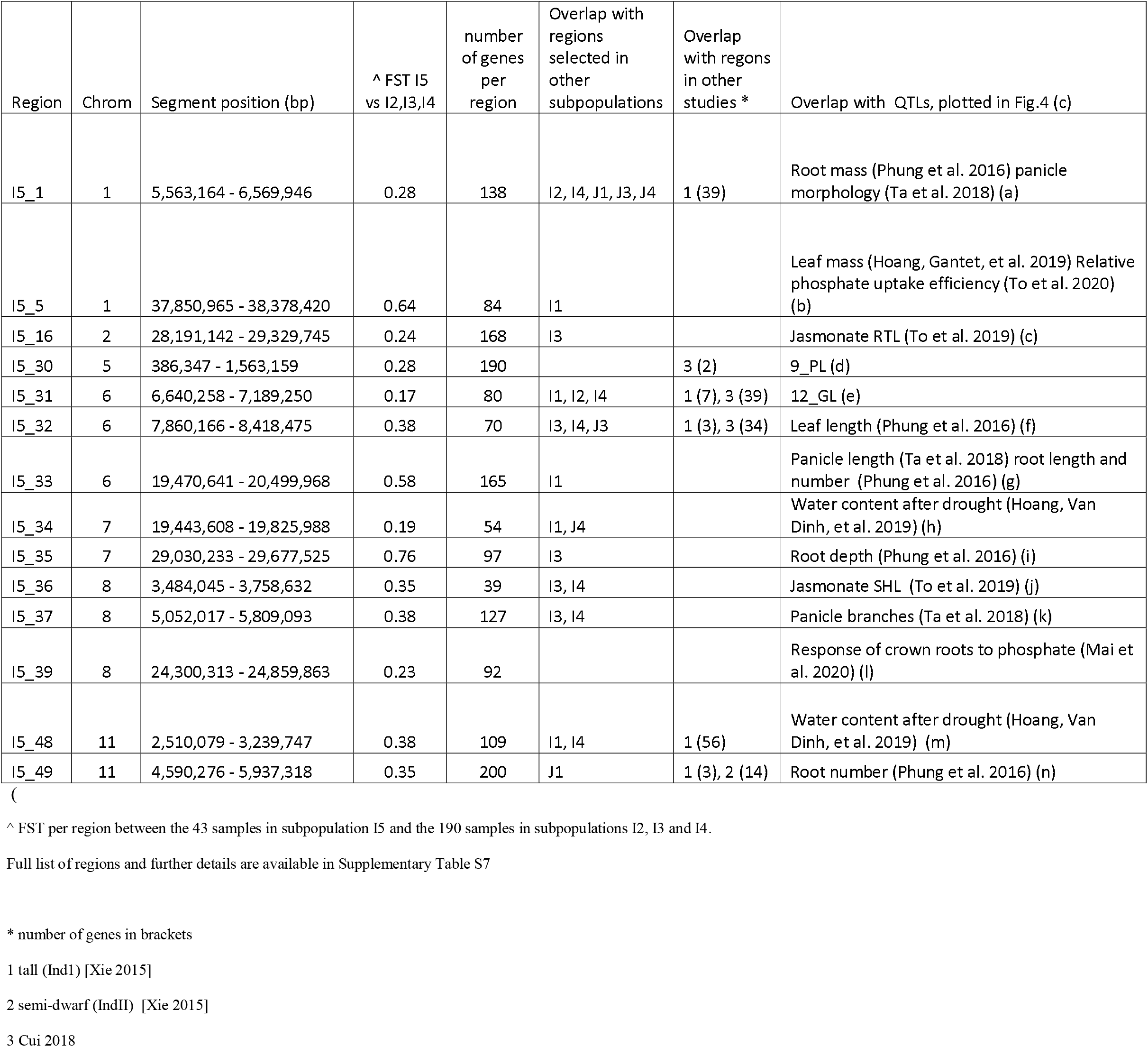
14 of the 52 regions under selection in the Indica I5 subpopulation, which overlap with QTLs. Detailing the overlap of selected regions with blished QTLs for Vietnamese rice populations, selected regions in Indica and Japonica subpopulations, and published selected regions (Xie et al. 2015; Lyu l. 2014; Cui et al. 2019)

### Candidate genes and non-synonymous alleles in selected regions of I5

We carried out three further analysis on the I5 subpopulation selected regions; a functional enrichment of the genes found within the 52 regions (Supplementary Table S8), a detailed annotation of the 4,576 genes in the 52 regions selected in the I5 subpopulation (Supplementary Table S10), and an analysis of the location of twenty-one genes with a putative role in salt tolerance in rice (Ganie et al. 2019) within the regions selected in the I5 subpopulation (Supplementary Table S12). Eventually, sixty-five candidate genes among the genes identified in the previous three analysis were short-listed using the following criteria (Table 5, details in Supplementary Table S13); F_ST_ over 0.5 in the whole selected region or in the functionally enriched genes within regions, presence of “*High* impact” SNPs, and presence of candidate genes from overlapping QTL. Ten of the 65 genes contained SNPs classified using SnpEff as having a “*High* impact”, i.e. predicted to cause deleterious gene effects (five stop gains, two start loses and three frame swifts). The alleles of eight of the genes containing “High impact” SNPs were different in the I5 subpopulation compared to the other Indica subpopulations (Fig. 5 and Supplementary Table S14). Among these eight genes, five of them showed the same allele than the Japonica subpopulations. However, two genes (LOC_Os10g35604 and LOC_Os11g10070/OsSEU2) had alleles unique to the I5 subpopulation.

**Table 5:**
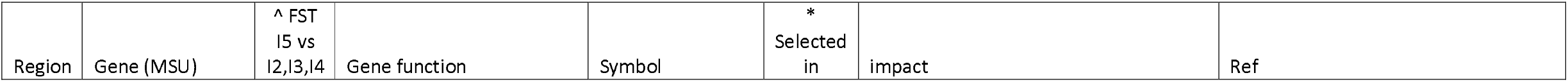
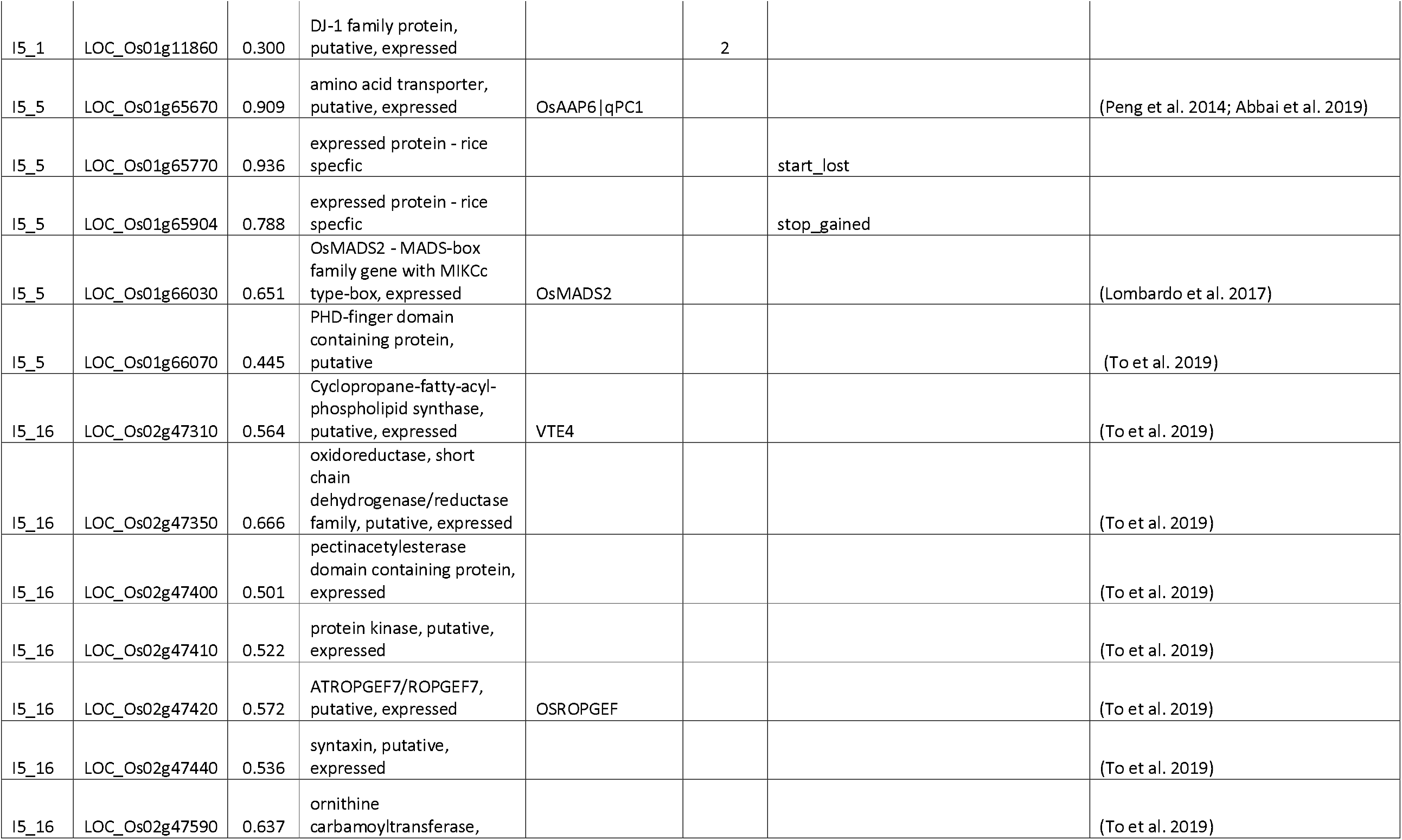
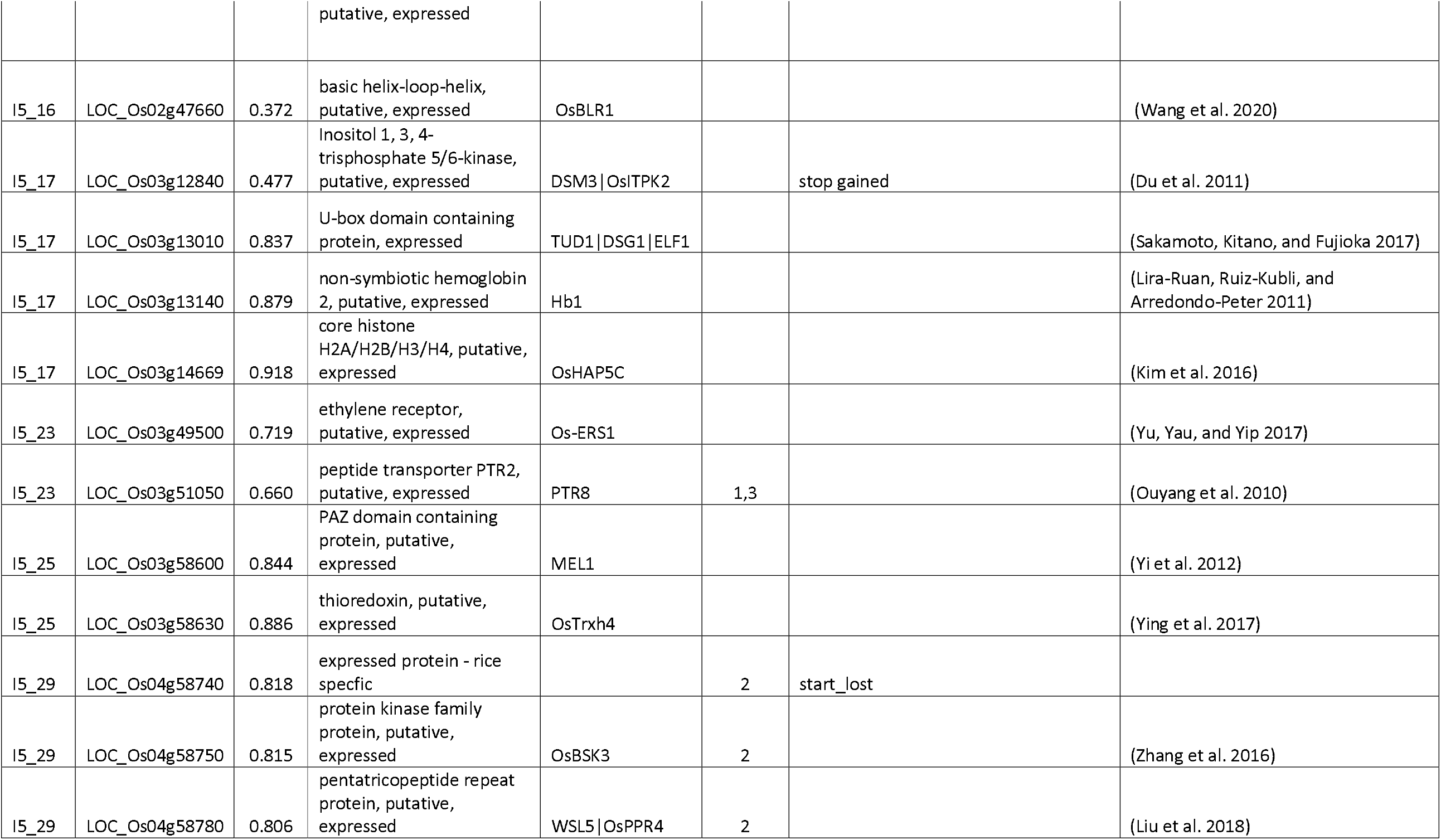
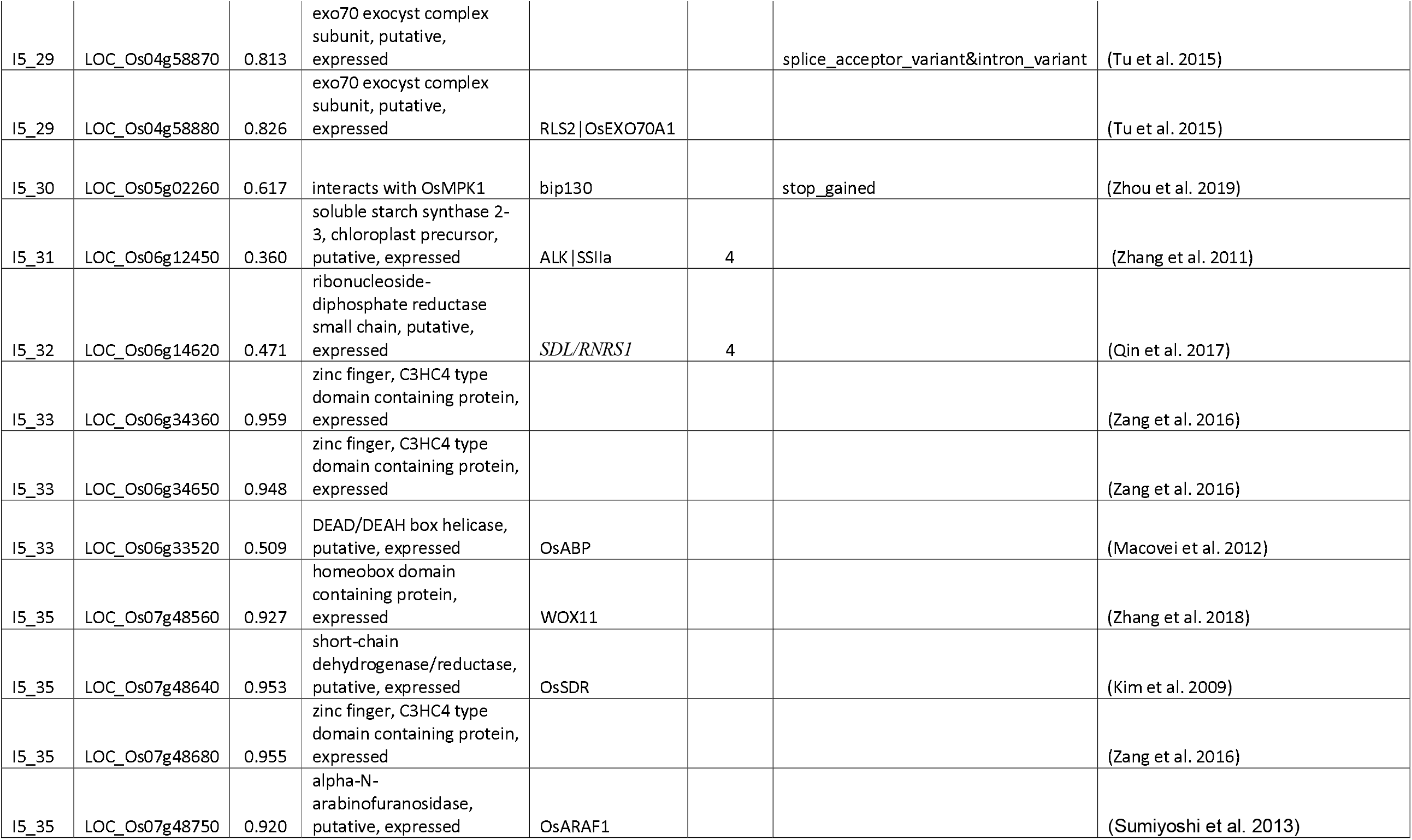
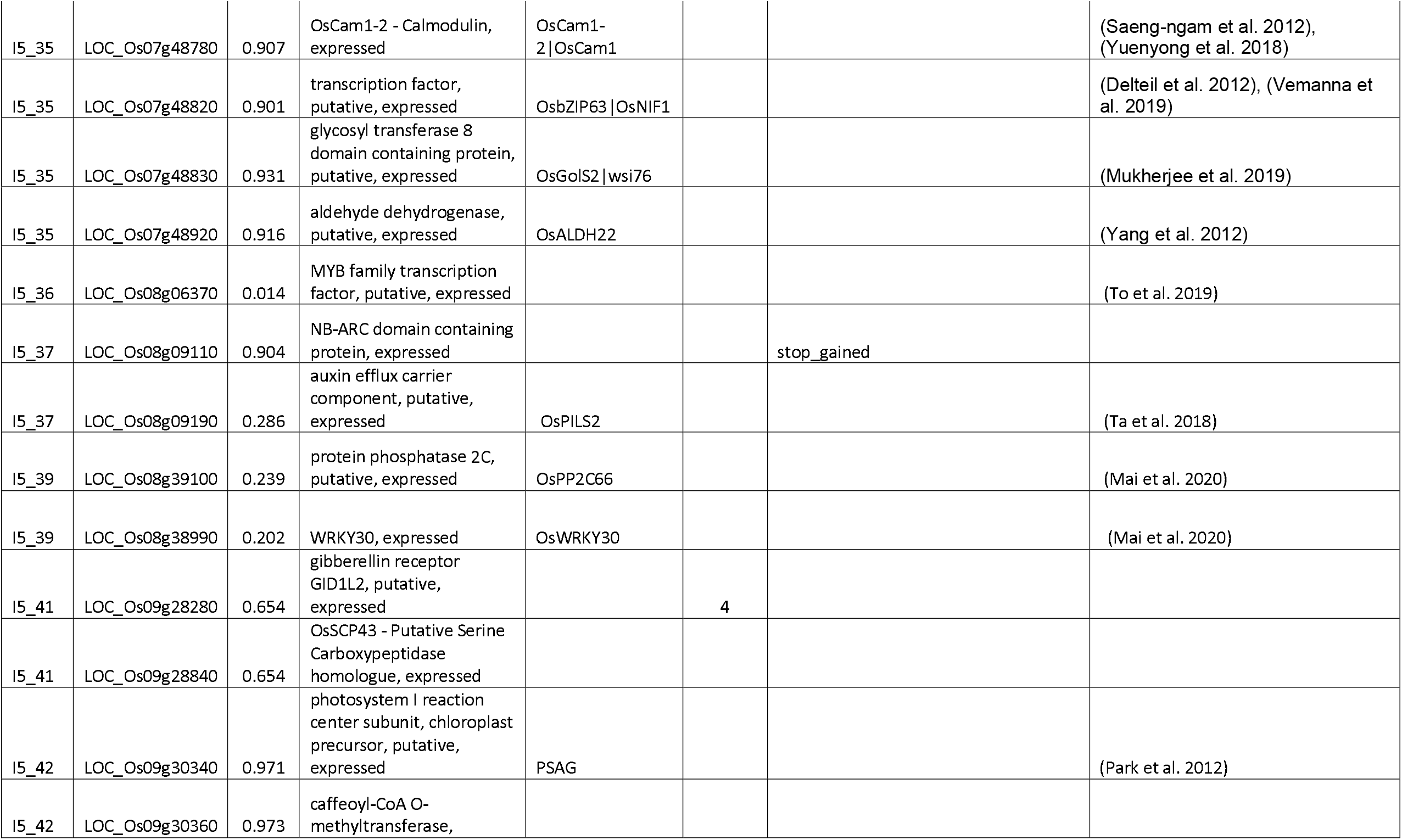
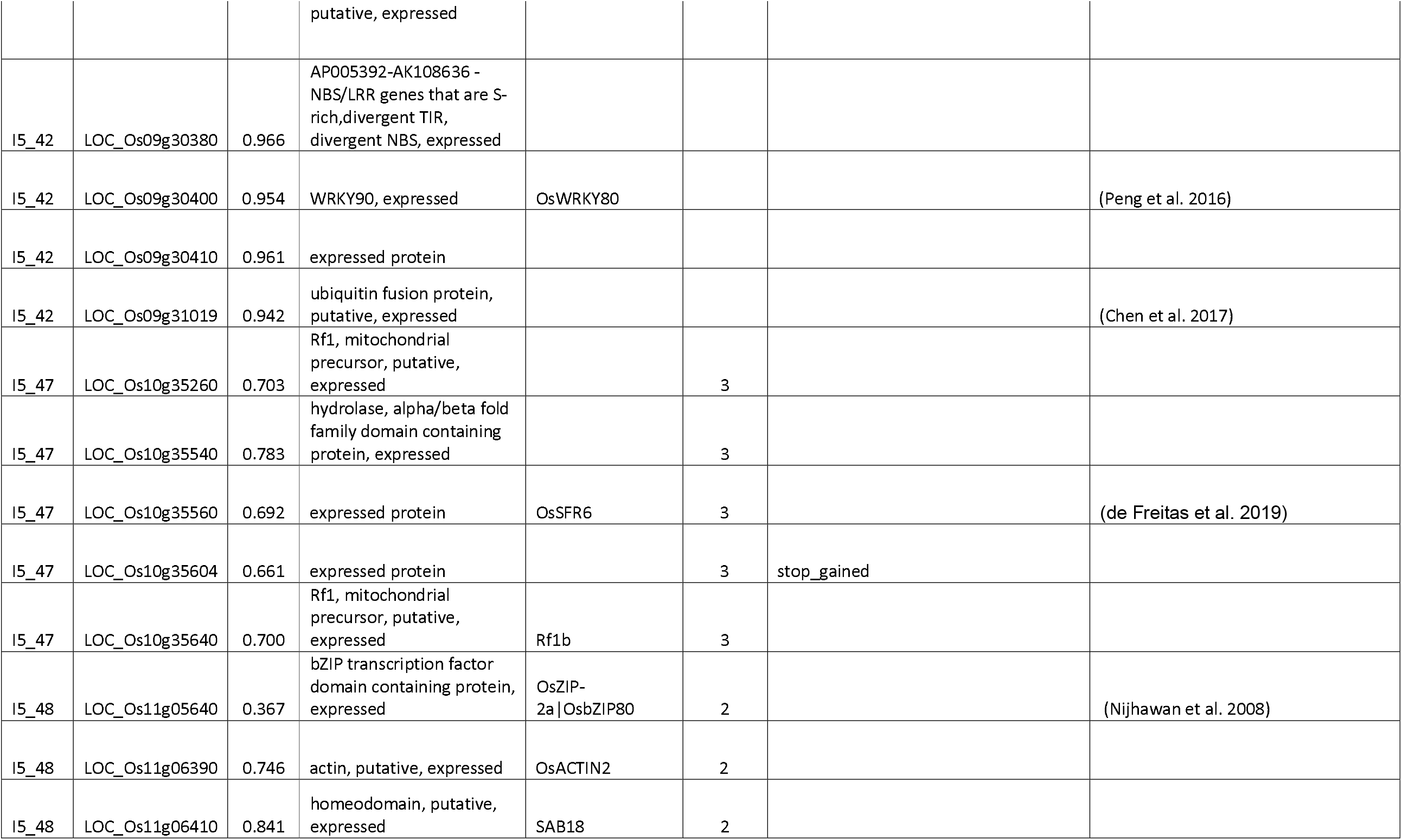
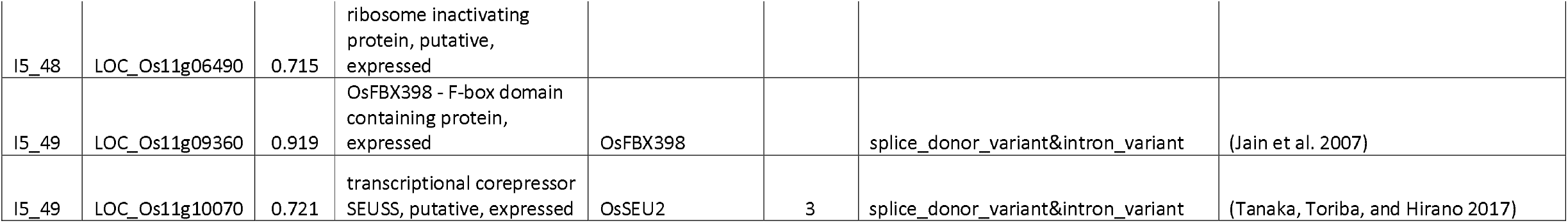
Candidate genes under selection in the Indica I5 subpopulation. Functional annotation of the 65 candidate genes and overlap with genes selected in previous dies *1. ecotype differentiated genes (Lyu et al. 2014)x, 2. tall (Ind1) (Xie et al. 2015)3. semi-dwarf (IndII) (Xie et al. 2015) 4. domestication(Cui et al. 2019) ^FST per gene between the 43 samples in subpopulation I5 and the 190 samples in subpopulations I2, I3 and I4. ther details are available in Supplementary Table S13

**Fig. 5.**
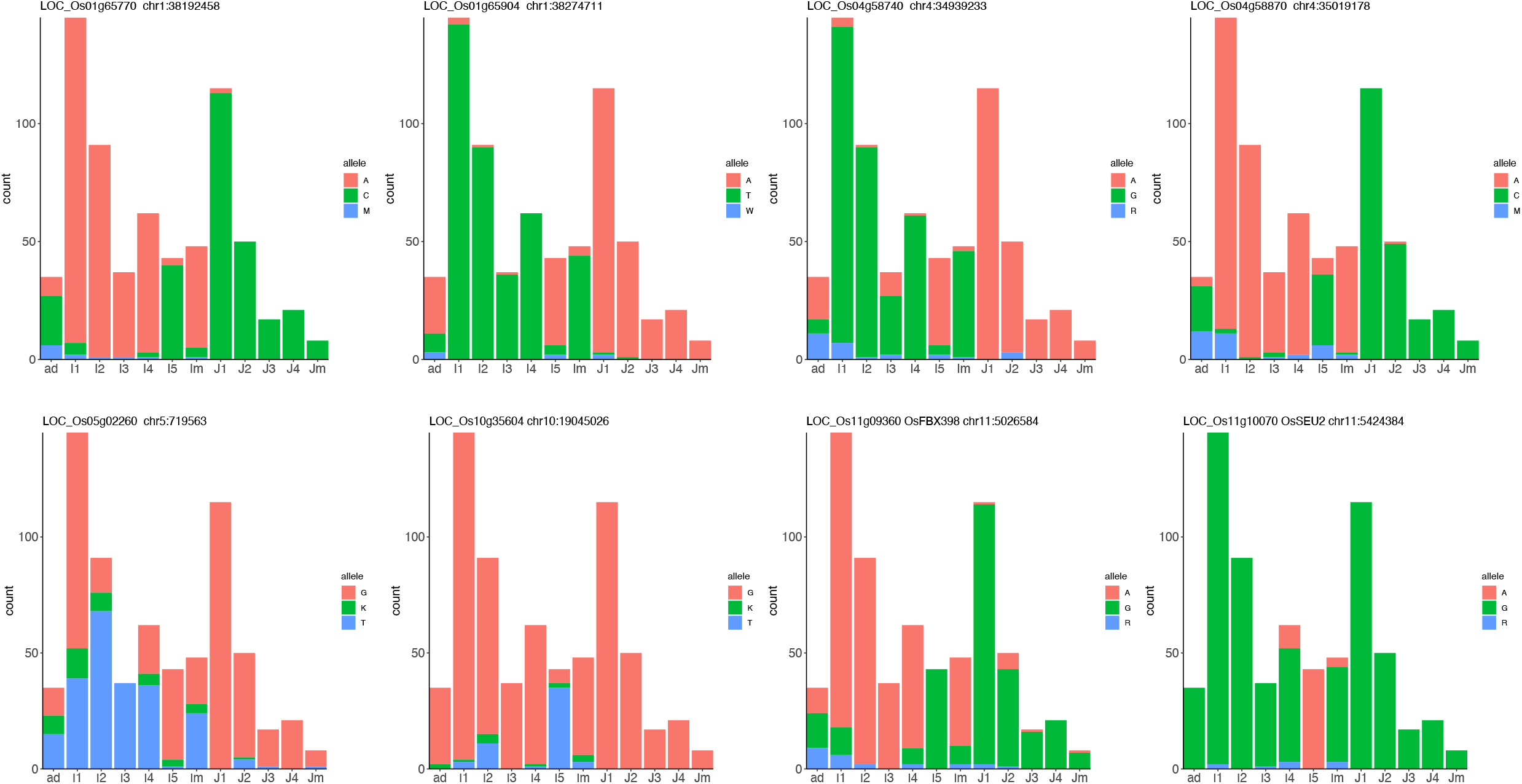
Allele Plots for “*High* impact” SNPs within eight candidate genes. Bar plots showing the base count for each subpopulation. A=adenine, T=thymine G = guanine, C = cytosine. Heterozygous calls are shown using IUPAC ambiguity codes.

## Discussion

Vietnam has one of the richest rice germplasm resources with over 4000 years of rice-cultivating experience. Local farmers have bred varieties to suit their ecosystem and regional culinary preferences. These conscious and unconscious selection processes have resulted in detectable changes in allele frequencies at selected sites and their flanking regions. We used a well-tested method, named XP-CLR, to identify distorted allele frequency patterns in contiguous SNP sites that cannot be explained by random drift. To identify regions under selection, we leveraged the strong population structure recently described in Vietnam (Higgins et al. 2021), which comprised five Indica and four Japonica subpopulations of native rice accessions adapted to variable geography and latitude range. We observed a stronger signature of selection in the Indica subtypes than in the Japonica subtypes, which may reflect the higher diversity within the Indica subtypes in Vietnam. Taking into consideration the size and diversity in each subpopulation (Higgins et al. 2021), the whole-genome XP-CLR score was lower in the larger subpopulations (I1 and J1) and the subpopulations with the lower diversity.

However, this trend was not true in the subpopulation indica-5 (I5), which showed a higher selection score than the other subpopulations with comparable size and diversity. Within the Indica subtypes, the subpopulation I5 showed the highest XP-CLR score against the subpopulation I1, which supports a strong signature for selection in I5 compared to the modern bred varieties in I1. On the contrary, the lowest XP-CLR score was obtained when I5 was compared to the I3 subpopulation, which is adapted to upland ecosystems (Phung et al. 2014). This suggest I5 shares selection pressures and resilient traits with upland varieties. Intermediate XP-CLR scores were obtained for the comparison of I5 with the two lowland subpopulations I2 (Mekong Delta) and I4 (Red River Delta).

Diversity is reduced when regions are under selection, but the observed diversity depends on many factors, including how long ago the selection occurred and the type of alleles selected alongside. This is referred to as the hitchhiking effect (Pavlidis and Alachiotis 2017). The fixation index (F_ST_) is a measure of population differentiation due to genetic structure. Both measurements vary highly along the genome but can provide additional information about the selected regions identified using XP-CLR. In this study, we calculated F_ST_ by comparing the I5 accessions to accessions in subpopulations I2, I3 and I4. We did not include the accessions in the elite I1 subpopulation, as we are specifically interested in genes that have been selected during the breeding of landraces within Vietnam. We used F_ST_ as an additional measure for identifying regions and genes under strong selection in the I5 subpopulation, and in support of the selection measurements obtained using XP-CLR.

Assigning functional roles to both regions and genes within the regions was the following natural step to identify breeding targets. We used two approaches, overlap with QTLs and functional enrichment. Seven QTL studies have been carried out on this dataset, finding associations for a range of traits relating to yield, this enables us to propose functional associations for around a third of the selected regions. A functional enrichment analysis evidenced selected regions were more clearly associated to specific GO terms in the Indica subpopulations than in the Japonica ones. The enrichment of GO terms was not correlated with the total number of genes or genome length in each subpopulation.

Looking in more detail at the 52 regions selected in the I5 subpopulation using a range of criteria, we identified 65 candidate genes within 20 of the selected regions. Six of these regions had a mean F_ST_ over 0.5 and we highlighted the following candidate genes within these regions. In region I5_35, we identified the transcription factor *WOX11* involved in crown root development (Zhang et al. 2018) and *OsCam1, OsbZIP63*, and *OsSDR*, which have putative roles in defence (Kim et al. 2009). Further genes of interest were (i) *OsAAP6*, a regulator of grain protein content (Peng et al. 2014), in region I5_5, (ii) *OsBSK3* (Zhang et al. 2016) and *WSL5* (Liu et al. 2018), which play roles in growth, in region I5_29, (iii) *OsABP*, which is upregulated in response to multiple abiotic stress treatments (Macovei et al. 2012), falls within region I5_33; and (iv) *OsSFR6*, a cold-responsive gene (de Freitas et al. 2019), in region I5_47. In addition, eight of the ten genes containing “*high* impact” mutations showed a different allelic content in the I5 subpopulation compared to the other Indica subpopulations, and in six cases these alleles were similar to the Japonica ones. Two genes containing “*high* impact” mutations were *OsFBX398*, an F-box gene with a potential role in both abiotic and biotic stresses (Jain et al. 2007; Vemanna et al. 2019), in region I5_49; and *bip130* (Zhou et al. 2019) in region I5_30, which regulates abscisic acid-induced antioxidant defence and fall within our QTL for panicle length (9_PL). In order to pinpoint candidate genes for a range of agronomic traits, we looked for overlap of selected regions with relevant QTLs. 14 of the 52 regions selected in the I5 subpopulation had overlaps with a wide range of QTLs, two of most relevant genes in these regions were *SSIIa*, which is responsible for the eating quality of rice (Zhang et al. 2011), and *OsbZIP80*, which is a transcription factor involved in dehydration stress response (Nijhawan et al. 2008).

Finally, we looked for overlaps with selected genes identified in three published studies using XP-CLR in rice (Lyu et al. 2014; Xie et al. 2015; Cui et al. 2019). Lyu et al. (2016) identified 56 Indica-specific genes in selected regions, which may account for the phenotypic and physiological differences between upland and irrigated rice. Thirty-one of these genes were on chromosome 3 and lied within regions also selected in the I4 and I5 subpopulations (I5_23, I5_24). The gene with the highest F_ST_ (0.67) is *ptr8* (LOC_Os03g51050), which encodes a peptide transporter (Ouyang et al. 2010). Xie et al. (2015) identified 2,125 and 2,098 coding genes in regions selected in the Chinese landraces (IndI) and modern-bred (IndII) subpopulations, respectively. We evidenced an overlap of 131 genes in selected regions in the I5 subpopulation with the genes selected in the IndI subpopulation and an overlap of 235 genes with the genes selected in the IndII subpopulation. This includes 7 genes in I5_22 and two genes in I5_23, both regions on chromosome 3, which were selected in all three subpopulations. Cui et al (2018) identified 186 potential selective-sweep regions in the Indica subtypes, of which 33 overlap with 9 of the 52 regions identified in the I5 subpopulation. These 9 regions contained 153 genes (Table 2). Cui et al. were specifically addressing the role of indigenous farmers in shaping the population structure of rice landraces in China, there is the possibility that similar regions may also have been selected in Vietnam. Substantial overlaps were found in three regions. On chromosome 2, 3 regions overlapped with I5_14. On chromosome 6, 11 regions overlapped with I5_31 and I5_32, including gene SIIa (LOC_Os06g12450), which is an important agronomic gene which is responsible for the eating quality of rice and plays an important role in grain synthesis. On chromosome 9, 13 regions overlapped with I5_4, including gene LOC_Os09g28280, which is a putative gibberellin receptor GID1L2 detailed in Table 2.

XP-CLR has proved a valuable method for identifying regions selected in the Vietnamese rice subpopulations and provided an insight into how natural selection and agricultural practices of farmers in Vietnam have shaped the population structure. Annotation of these regions with both overlaps with QTLs for a range of agronomic traits and functional enrichment allowed us to prioritise candidate regions as targets for breeding programs. Our results give further support for the Indica I5 subpopulation being an important source of novel alleles for both national and international breeding programmes. Using a range of criteria, F_ST_ and diversity in these regions, we identified 65 genes which could be further investigated for their breeding potential.

## Acknowledgements

The author(s) acknowledge the support of the Biotechnology and Biological Sciences Research Council (BBSRC), part of UK Research and Innovation; this research was funded by the BBSRC Core Strategic Programme Grant (Genomes to Food Security) BB/CSP1720/1 and its constituent work package BBS/E/T/000PR9818 (WP1 Signatures of Domestication and Adaptation); and BBSRC’s grants BB/N013735/1 (Newton Fund), and the Newton Fund Institutional Links (Project 172732508), which is managed by the British Council.

## Author contribution statement

TDK, KHT, AH, SD, LHH, BS, MC and JDV designed and conceived the research. JH performed the data analysis with assistance from JDV. JH and JDV wrote the paper. All authors read and approved the final manuscript.

## Availability of data

All sequence data used in this manuscript have been deposited as study PRJEB36631 in the European Nucleotide Archive.

## Supplementary tables

Supplementary Table S1 Reciprocal difference between comparisons for each subpopulation pair

Supplementary Table S2. Summary of XP-CLR comparisons for the Indica and Japonica subpopulations. Detailing the XP-CLR mean, cut off and number of regions for each comparison.

Supplementary Table S3. List of genes selected in each subpopulation.

Supplementary Table S4. Overlap of the selected regions in Indica subpopulations with QTL found in Vietnamese rice datasets.

Supplementary Table S5. Overlap of the selected regions in Japonica subpopulations with QTL found in Vietnamese rice datasets.

Supplementary Table S6. Gene Ontology enrichment for each selected region.

Supplementary Table S7. List of 52 regions selected in the Indica I5 subpopulation.

Supplementary Table S8. Annotation of I5 selected regions using PhytoMine. Enrichment analysis for protein domain and Meta-Cyc pathway using PhytoMine.

Supplementary Table S9. List of genes selected in each region for the Indica I5 subpopulation.

Supplementary Table S10. List of all genes selected in I5 subpopulation. Detailing MSU and RAP gene ID, annotation and enrichment in Phytomine, high impact SNPs and mean F^st^

Supplementary Table S11. Details of QTLs within the fourteen overlapping regions.

Supplementary Table S12. List of 21 genes related to salt tolerance selected in the I5 subpopulation.

Supplementary Table S13. Details of 64 candidate genes.

Supplementary Table S14. Details of “*High* impact” SNPs within eight candidate genes.

## Supplementary Figures

Fig. S1 Chromosome plots of regions selected in each Indica subpopulation showing the regions selected against each individual subpopulation and the shaded final selected regions which were selected against three subpopulations. a) 44 regions selected in I1, b) 41 regions selected in I2, c) 42 regions selected in I3, d) 38 regions selected in I4, e) 52 regions selected in I5

Fig. S2 Chromosome plots of regions selected in each Japonica subpopulation showing the regions selected against each individual subpopulation and the shaded final selected regions which were selected against two subpopulations. a) 28 regions selected in J1, b) 23 regions selected in J2, c) 24 regions selected in J3, d) 25 regions selected in J4

Fig. S3 Range of sizes for selected regions in (a) five Indica subpopulations and (b) four Japonica subpopulations

Fig. S4 Upset plots for overlap of genes in selected regions for (a) all nine subpopulations, (b) five Indica subpopulations and (c) four Japonica subpopulations

