## Supplementary figure S1 for "Identifying genomic regions and candidate genes selected during the breeding of rice in Vietnam"

**Figure S1. Chromosome plots of regions selected in each Indica subpopulation showing the regions selected against each individual subpopulation and the shaded final selected regions which were selected against three subpopulations.**

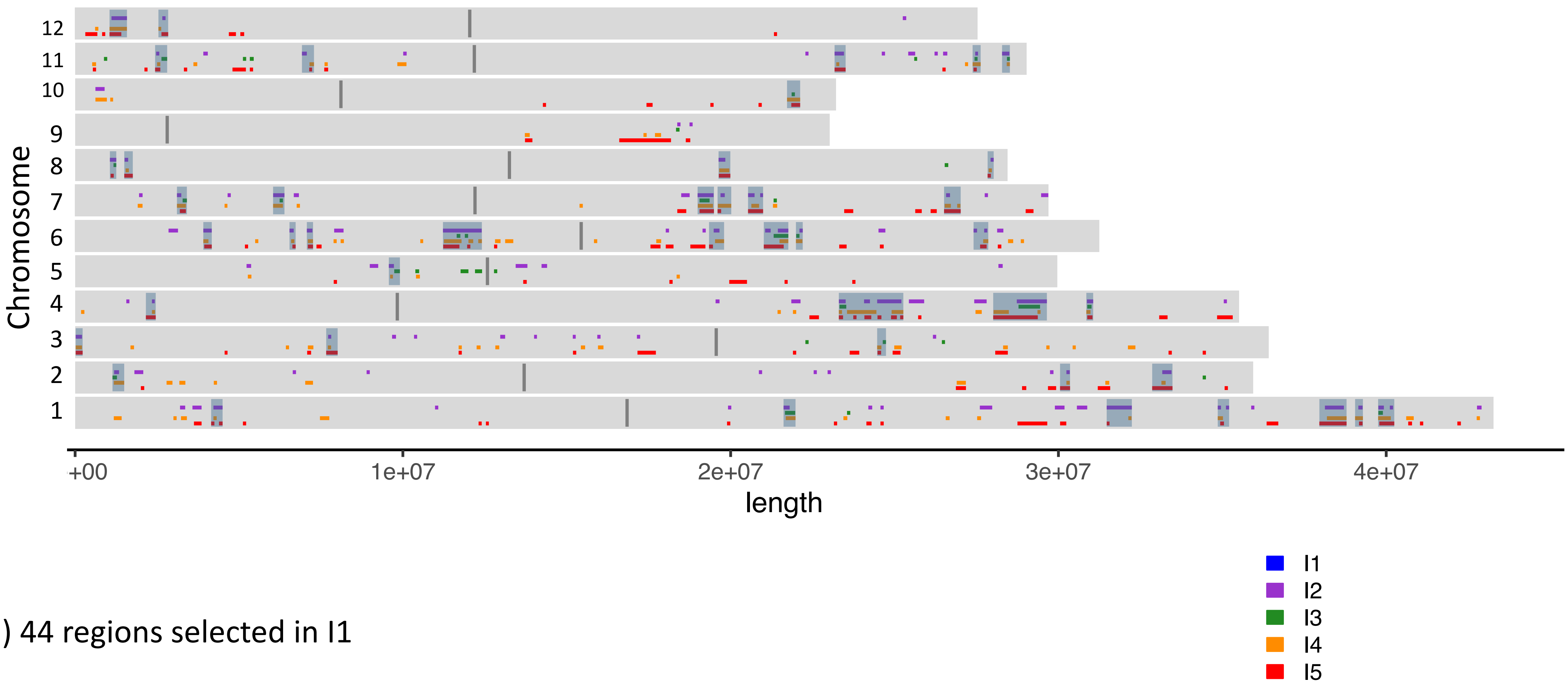


a) 44 regions selected in I1

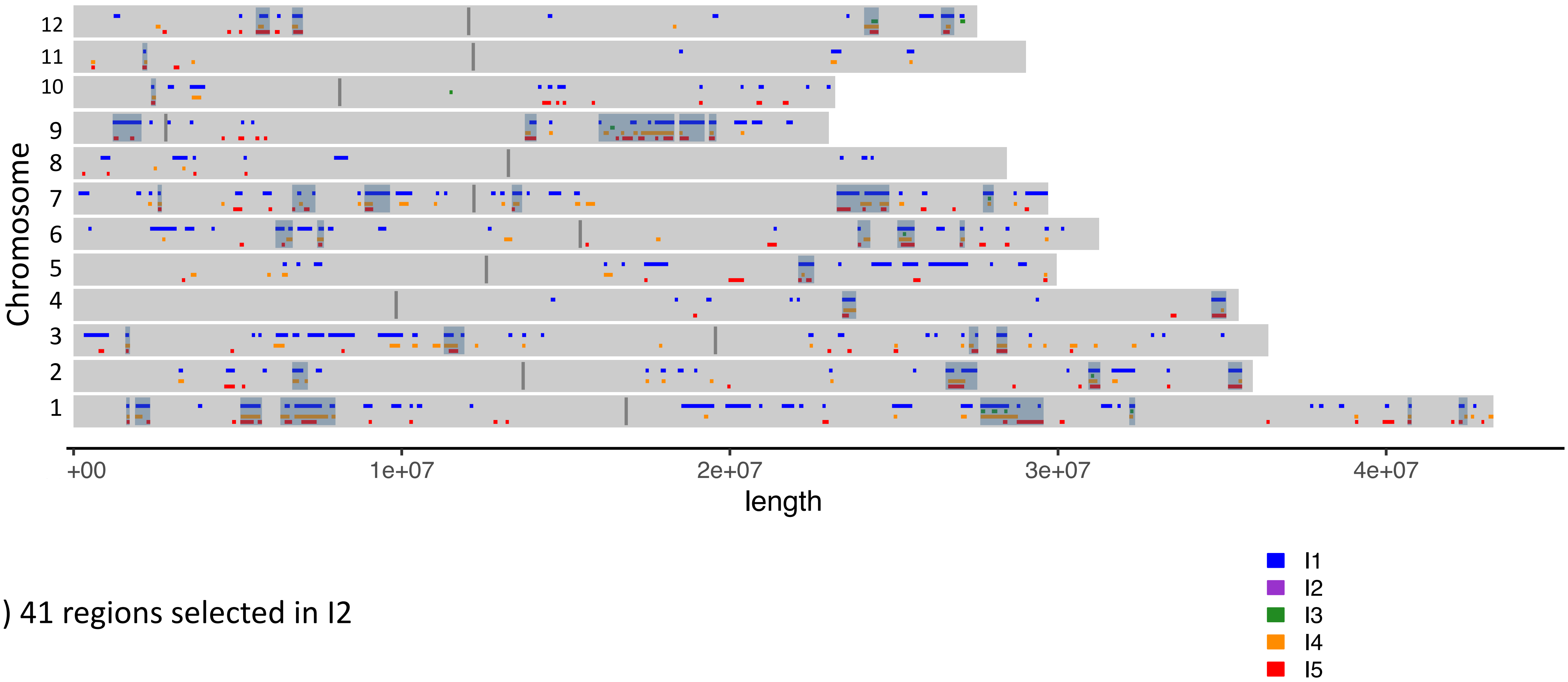


b) 41 regions selected in I2

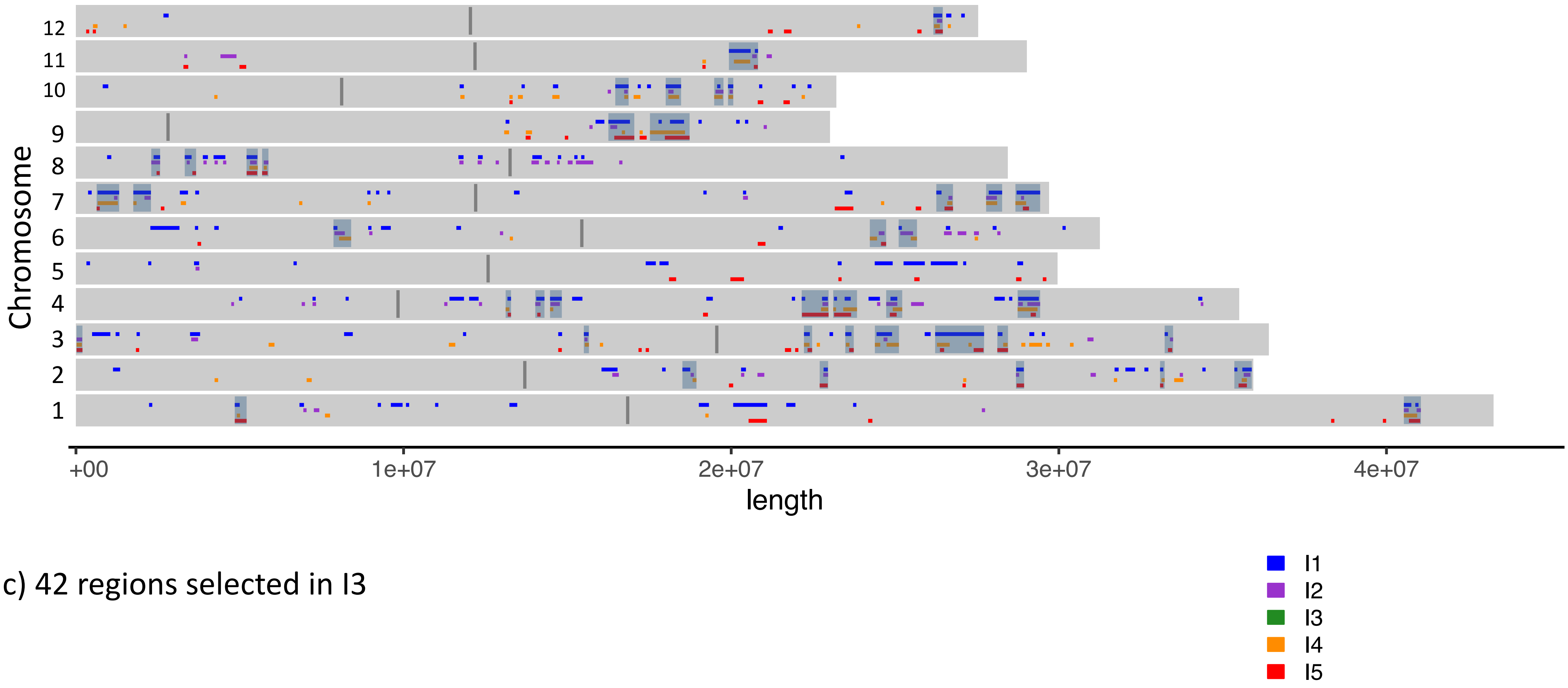


c) 42 regions selected in I3

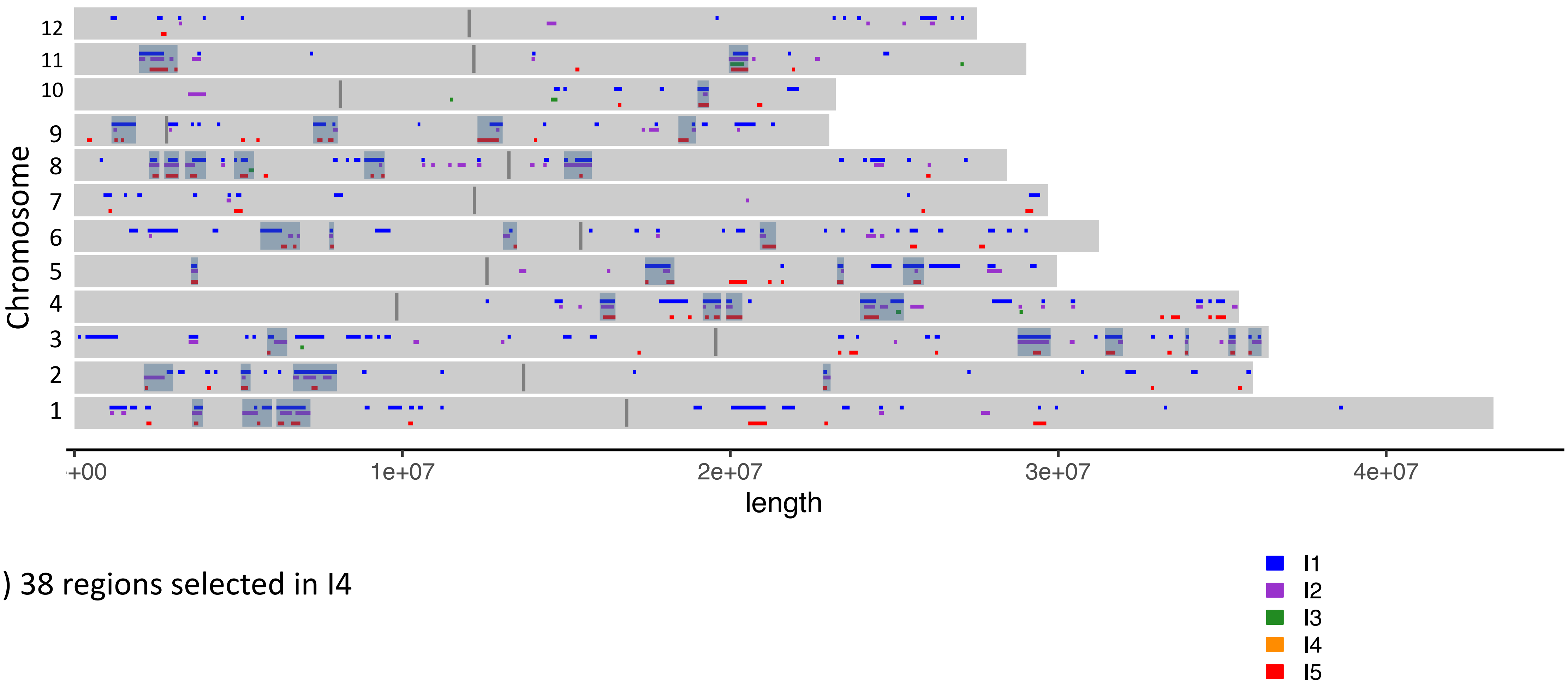


d) 38 regions selected in I4

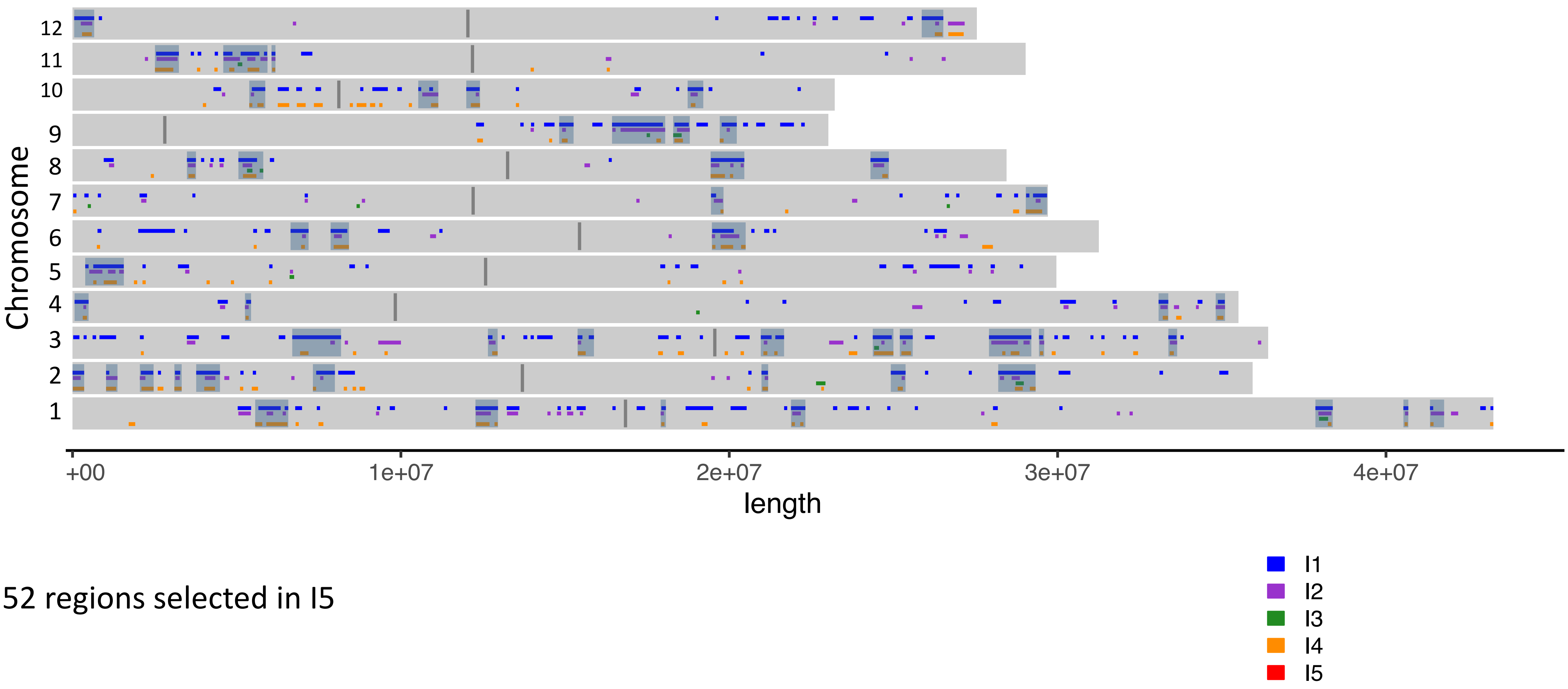


e) 52 regions selected in I5
