## Supplementary figure S2 for "Identifying genomic regions and candidate genes selected during the breeding of rice in Vietnam"

**Figure S2. Chromosome plots of regions selected in each Japonica subpopulation showing the regions selected against each individual subpopulation and the shaded final selected regions which were selected against two subpopulations.**

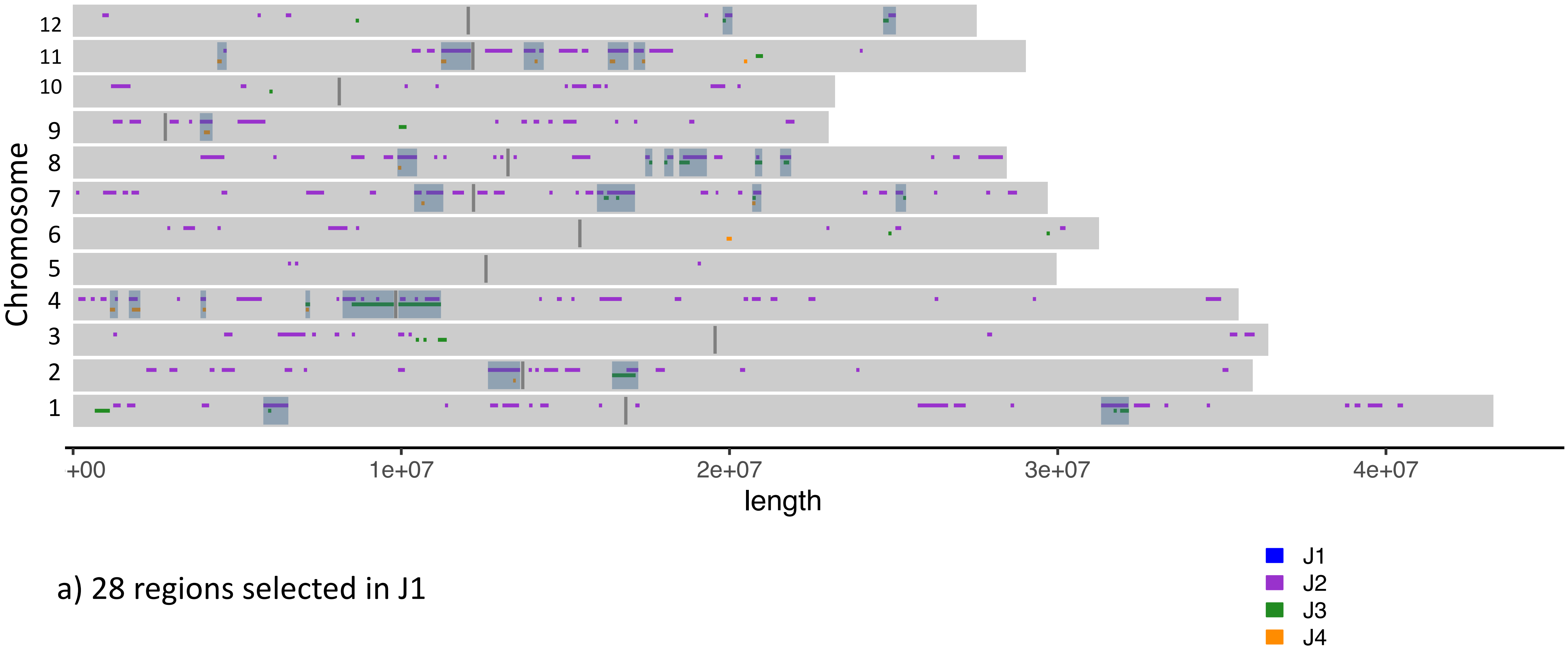


a) 28 regions selected in J1

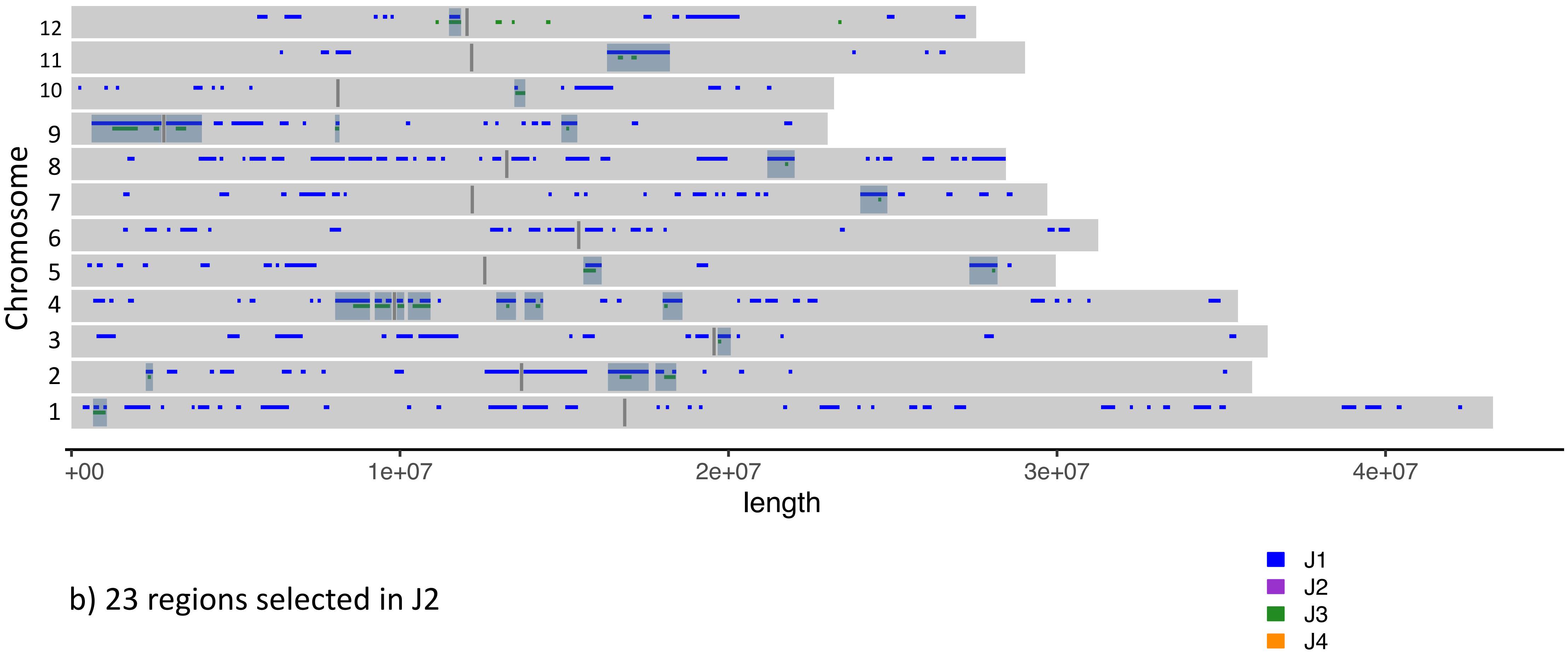


b) 23 regions selected in J2

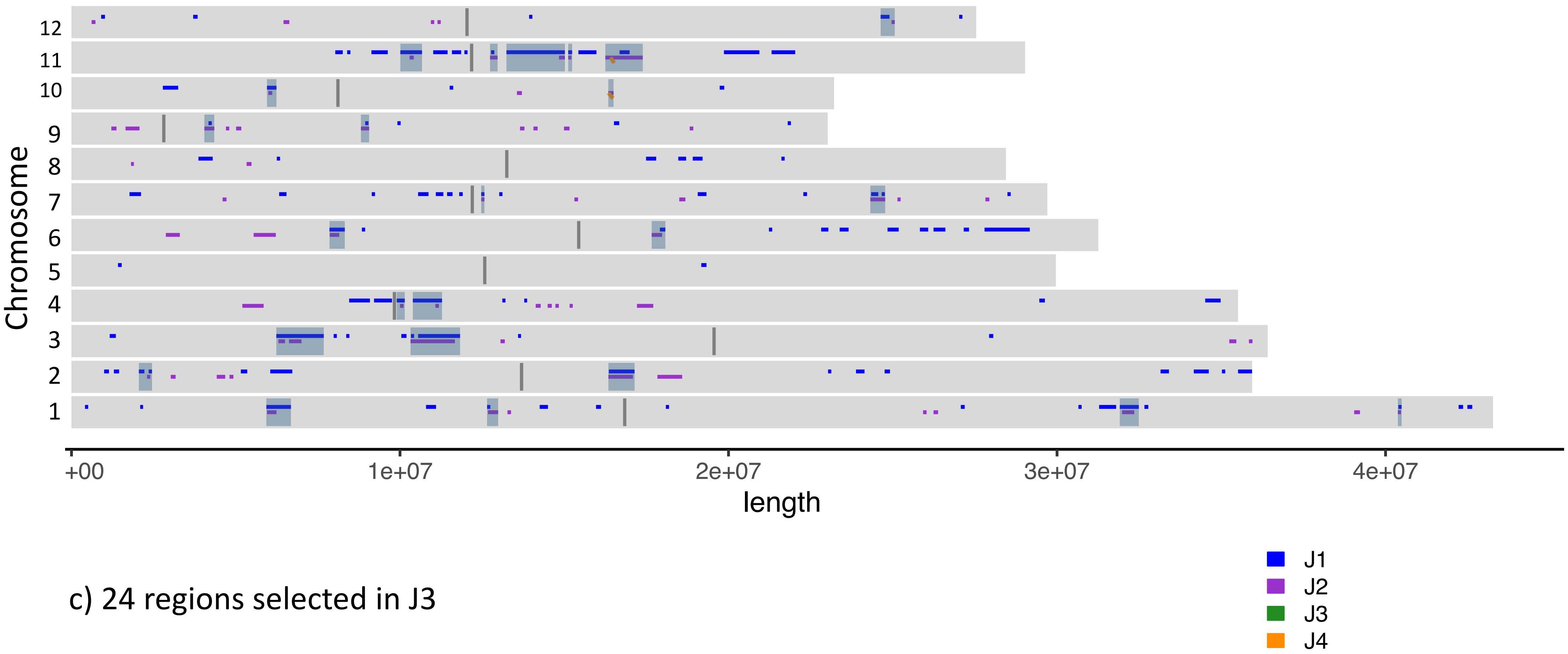


c) 24 regions selected in J3

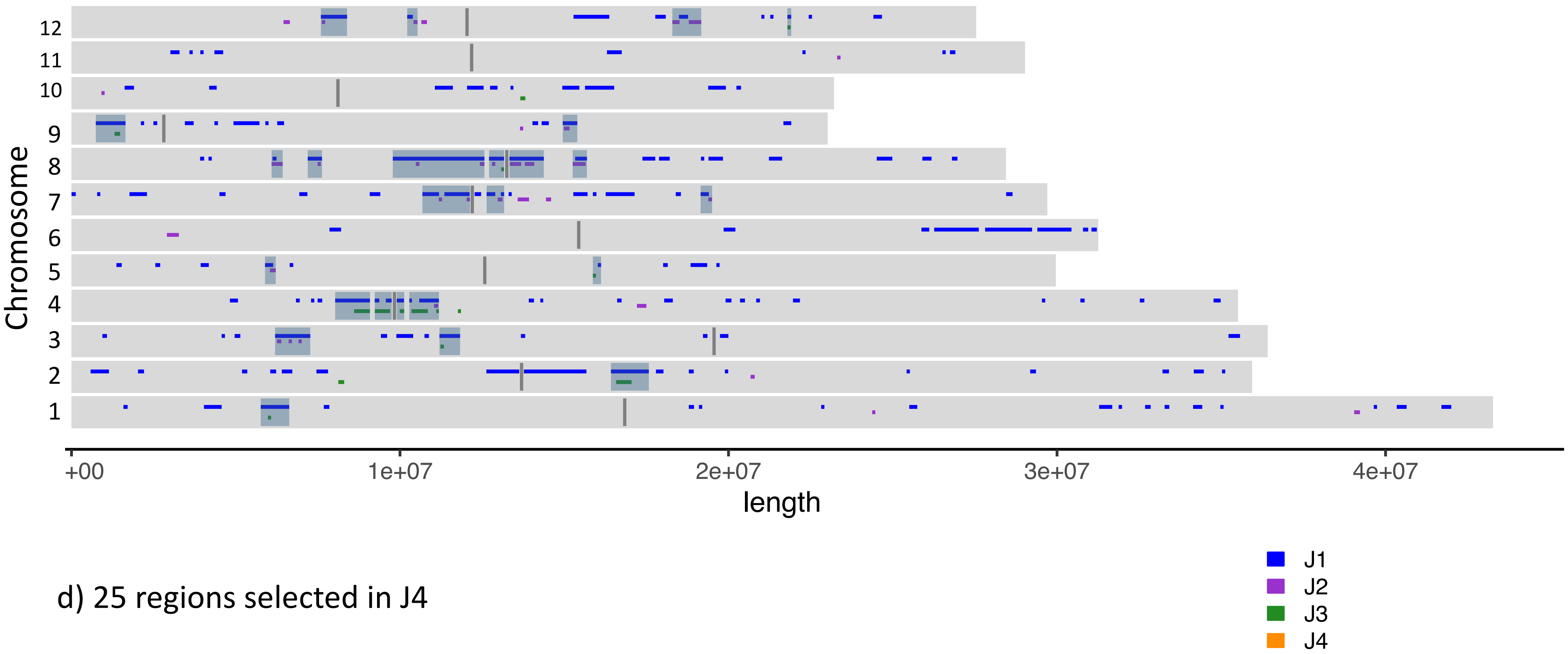


d) 25 regions selected in J4
