## Supplementary figure S3 for "Identifying genomic regions and candidate genes selected during the breeding of rice in Vietnam"

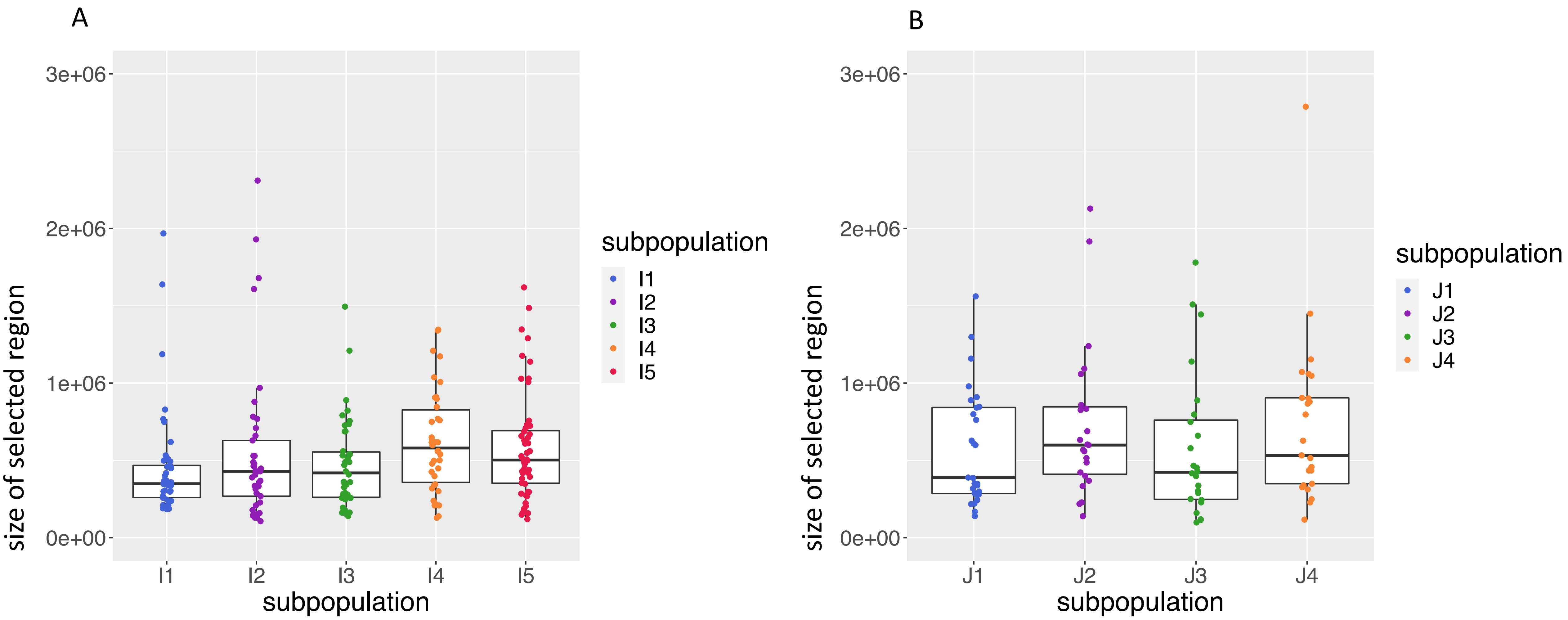


Fig. S3 Range of sizes for selected regions in (a) five Indica subpopulations and

(b) four Japonica subpopulations
