## Supplementary figure S4 for "Identifying genomic regions and candidate genes selected during the breeding of rice in Vietnam"

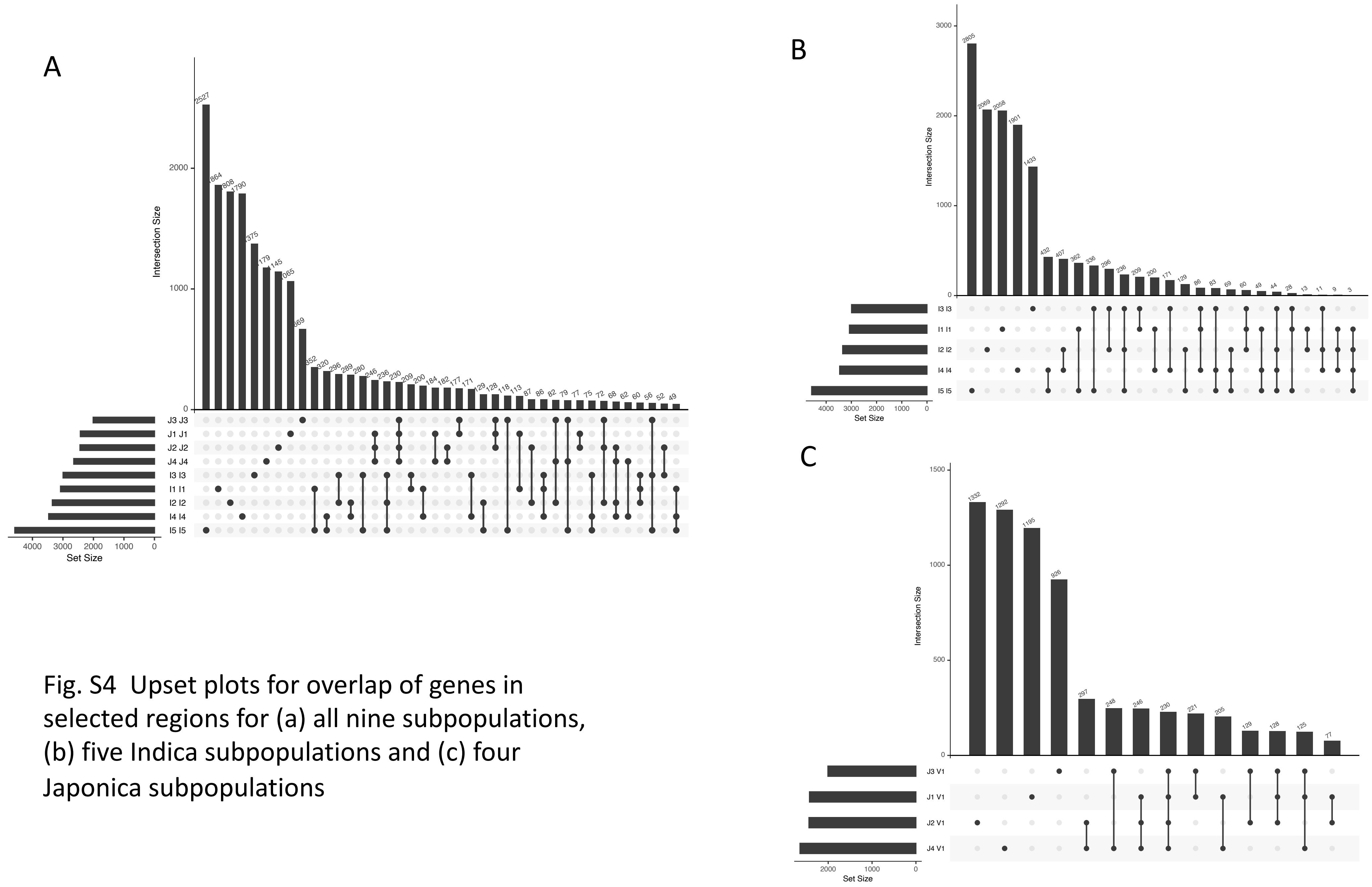


Fig. S4 Upset plots for overlap of genes in selected regions for (a) all nine subpopulations, (b) five Indica subpopulations and (c) four Japonica subpopulations
